# Sexually dimorphic plasticity of PV inhibition in sensory neocortex during learning

**DOI:** 10.64898/2026.01.10.698814

**Authors:** Eunsol Park, Dika A. Kuljis, Stephanie E. Myal, Joseph A. Christian, Alison L. Barth

## Abstract

Neocortical parvalbumin-expressing (PV) neurons critically regulate circuit excitation by strong synaptic inputs onto local pyramidal (Pyr) neurons. Plasticity in PV-mediated inhibition during learning could have pronounced effects on gating excitatory synaptic plasticity and circuit excitability, but experimental evidence to support this input- and target-specific plasticity is scant. Here, we combined *in vitro* electrophysiology with quantitative synapse analysis to determine whether training in a whisker-based sensory-association task could alter PV-mediated inhibition in the primary somatosensory cortex of mice. Using light-evoked activation of channelrhodopsin-expressing PV neurons, we found that evoked PV-IPSCs in Pyr neurons from layer (L) 2/3, but not L5, were rapidly suppressed at the onset of training. This reduction was sex-specific, occurring only in females. The training-related decrease in PV output was accompanied by a reduced number of PV-associated synapses on both the soma and dendrites of L2/3 Pyr neurons, suggesting a postsynaptic structural change. Notably, when whisker stimulation was decoupled from the water reward during pseudotraining, PV-mediated inhibition remained stable. Thus, reduced PV inhibition in superficial layers is an early response to the development of stimulus-reward associations during sensory learning. In addition, these data underscore the importance of including sex as a biological variable in studies of learning-related cortical plasticity.

## Introduction

Reducing inhibition has long been appreciated as an important mechanism for gating excitatory synaptic plasticity and circuit rewiring that occur during experience and learning^1–3^. Disinhibition of pyramidal (Pyr) neurons can be controlled directly via the suppression of parvalbumin (PV) or somatostatin (SST)-expressing interneurons that directly innervate Pyr neurons. Alternatively, disinhibition can be mediated indirectly, by the activation of other GABAergic neurons that specifically target PV and SST neurons^4–8^. Although disinhibition can be short-term and reversible, initiated by transient changes in circuit activity or brain state^4,6,7,9,10^, long-lasting changes in inhibition have also been associated with learning, particularly at the early stages^11–14^.

Learning-related changes in both the activity and synaptic output of SST neurons have been identified^15–19^; however, how learning alters the activity or synaptic properties of PV neurons is less clear. Anatomical analysis indicates that contextual fear conditioning can alter both PV expression and bouton number in the hippocampus^13^, and acquisition of a skilled motor learning task transiently increases PV boutons in the superficial layers of motor cortex^14^. Longitudinal Ca^++^ imaging of PV neurons in visual cortex suggests that PV neurons become recruited into stimulus-specific ensembles during learning^20^. In prefrontal cortex, a subset of PV neurons shows increased gamma synchrony during rule-shift learning, a process required for successful behavioral adaptation^21,22^. These findings highlight dynamic changes in PV activity, but do not address whether there are long-lasting changes in inhibition from PV neurons, or whether PV inhibition across the cortical column is differentially affected. Notably, PV output depression may contribute to critical period plasticity in auditory cortex^23^, and a reduction in PV neuron activity can enhance stimulus-evoked Pyr neuron responses in behaving animals^24,25^. Based upon these findings, we hypothesized that PV output plasticity may be involved in neocortical circuit changes during sensory learning.

We sought to determine whether PV to Pyr inhibition would be regulated in mouse somatosensory (barrel) cortex during association learning, where a multiwhisker (airpuff) stimulus predicted a subsequent water reward^26^. This training drives both *in vivo* response plasticity and also synaptic changes in selective neural subtypes across the cortical column^17,18,27–29^. Optogenetically-evoked PV output was evaluated in layer (L) 2/3 and L5 Pyr neurons in acute brain slices from control and trained mice, layers where previous studies have shown significant potentiation of thalamocortical and intracortical synaptic strength^27,28,30^. In addition, we used anatomical analysis of PV synapses onto the soma of Pyr neurons to identify learning-related changes in output. We identify a reduction in PV inhibition at the onset of learning, specific to conditions where the stimulus is predictive of reward delivery. Learning-related plasticity in PV-output was restricted to female mice and could not be attributed to reduced intrinsic excitability of PV neurons.

## Results

### PV synapse density on L2/3 Pyr neurons is reduced after sensory association training

To determine how PV-mediated inhibition onto Pyr neurons in the barrel cortex is modulated during associative learning, we trained freely moving PV-Cre transgenic mice in an automated homecage training environment^26,27^. Sensory association training (SAT) was initiated after 1-2 days of acclimation to the training cage (ACC), where mice can nose poke to initiate water release. During SAT, a gentle airpuff (∼6 psi, 500 ms) preceded water delivery (Fig. 1b). In 20% of initiated trials, neither airpuff nor water reward was delivered (blank trials). To evaluate whether animals associated the whisker stimulus with water delivery, we quantified the frequency of anticipatory licking for stimulus and no-stimulus, blank trials across the training period (see Materials and Methods). Consistent with previous studies^18,26,28,29^, approximately 70% of animals (28/42) showed greater licking frequencies for stimulus trials than for blank trials after 1 day of SAT (SAT1). Despite the heterogeneity of performance across animals, average licking frequencies for stimulus trials were significantly higher than for blank trials (Fig. 1h; Stim 5.1±0.3 Hz vs. Blank 4.3±0.3 Hz, p=0.0049). These results indicate that PV-Cre mice learned the stimulus-reward association at rates comparable to other transgenic strains^18,26–29^.

**Fig. 1.**
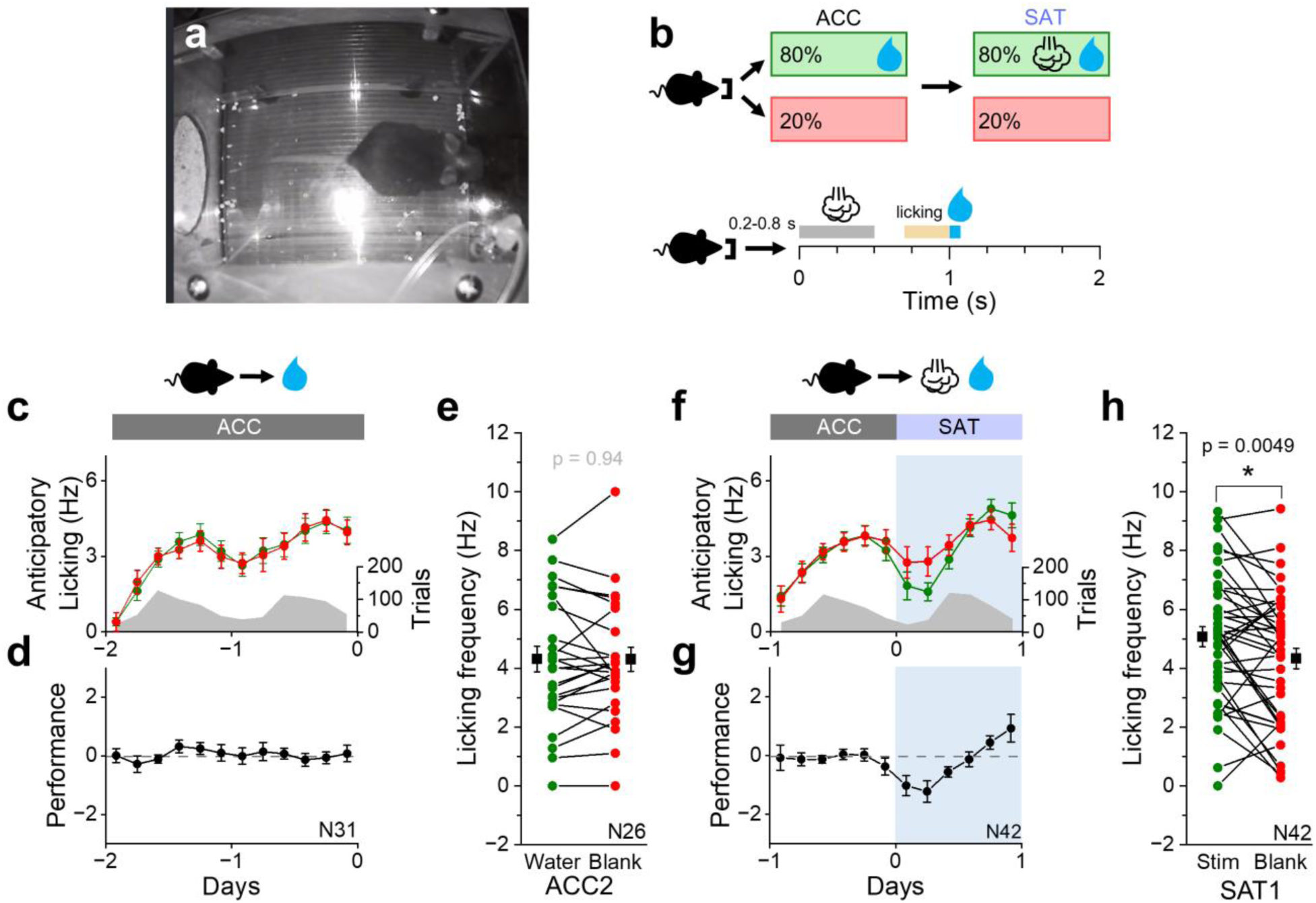
Mice form an association between whisker stimuli and water after sensory association training. (a) Schematic of the automated cage training set-up. (b) Top: Timeline of cage acclimation and sensory association training (SAT). SAT begins following a 1-2 day acclimation period. Bottom: Trial structure. Trials were initiated by a nosepoke, followed by a random delay (0.2-0.8 s), a gentle airpuff (500 ms, 6 psi), a fixed 500 ms delay, and then water delivery. Anticipatory licking frequency (Hz) was quantified during the 300-ms window preceding water delivery (yellow line) for both stimulus and blank trials. (c) Mean anticipatory licking frequency (Hz) for water (green) and blank (red) trials following 1-2 days of acclimation (ACC). Gray shading indicates the distribution of average trial numbers across time bins. Number of animals: 1-day ACC, N=5 mice; 2-day ACC, N=26 mice. (d) Performance (Lick_Water_-Lick_Blank_; see Materials and Methods) was calculated for each 4-hr bin and averaged across animals. (e) Mean anticipatory licking frequency (Hz) during the last 20% of water (green) and blank (red) trials for each animal after 2 days of ACC (ACC2). Black symbols represent group mean±SEM. Water: 4.3±0.4 Hz; Blank: 4.3±0.4 Hz; N=26 mice. Paired-sample, Wilcoxon signed-rank test (Z=-0.08, n=26). (f-h) Same as in (c)-(e), but for animals after 1 day of SAT. Blue shading indicates the training period. (h) Stim: 5.1±0.3 Hz; Blank: 4.3±0.3 Hz; N=42 mice. Wilcoxon signed-rank test (*Z*=2.76, n=42). * p<0.05.

After training animals, we performed fluorescence-based quantitative synapse analysis using the neuroligin-based synapse-tagging molecule, FAPpost^31,32^. As previously demonstrated, this method reliably detects PV synapses with high accuracy^31^. Pyr neurons were virally transduced with a cell-filling dTomato (dTom) and postsynaptic FAPpost in PV-Cre x Ai3 (Cre-dependent YFP) transgenic mice to label PV neurons with YFP comprehensively. Confocal imaging and digital alignment of presynaptic PV structures with postsynaptic sites tagged by FAPpost were used to examine the density of PV-assigned FAPpost puncta (putative PV synapses) on the soma and dendrites for a target Pyr neuron. FAPpost puncta were identified as PV synapses if they colocalized with PV neurites (see Materials and Methods).

Combining postsynaptic molecular markers with presynaptic neurite signals provides a robust method for detecting and quantifying input-specific synapses^31,33^. While PV neurons predominantly target Pyr neurons’ perisomatic regions, some innervation also extends to the dendrites^31^. To determine whether PV synaptic plasticity was compartment-specific, we separately quantified somatic and dendritic PV-associated synapse density. After SAT1, the density of PV-associated synapses at both the soma and dendrites decreased in L2/3 Pyr neurons (Fig. 2a-g; Soma: ACC 0.065±0.008/µm^2^ vs. SAT1 0.038±0.009/µm^2^, p=0.0052; Dendrites: ACC 0.29±0.04/µm vs. SAT1 0.18±0.03/µm, p=0.020). The decrease in number of PV-associated synapses did not depend on the density of presynaptic PV structures, because the PV neurite density along the soma and dendrites of Pyr neurons was not altered after SAT1 (Supplementary Fig. 1; Soma: ACC 0.19±0.02/µm^2^ vs. SAT1 0.17±0.02/µm^2^; p=0.43; Dendrites: ACC 1.4±0.2/µm vs. SAT1 1.2±0.1/µm; p=0.39).

**Fig. 2.**
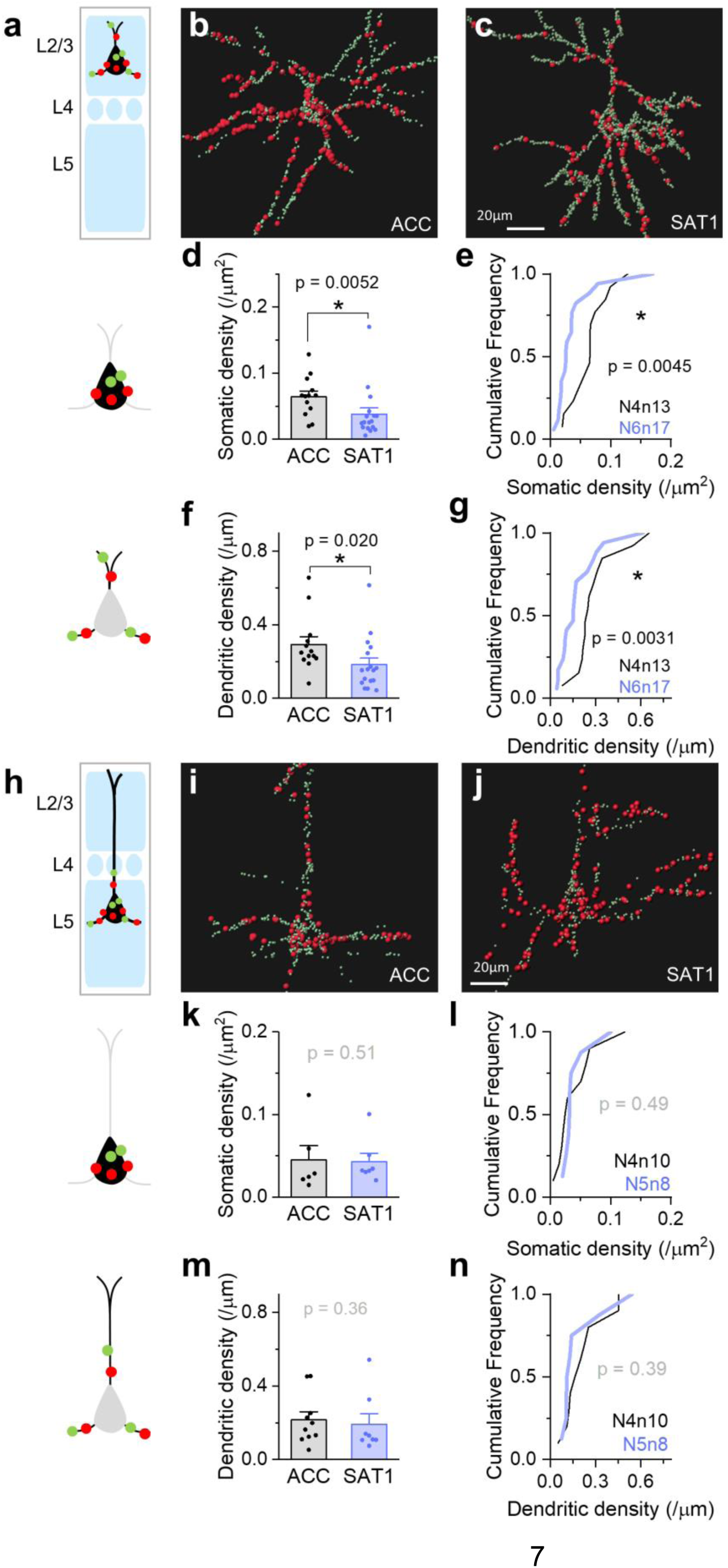
PV synapse density onto L2/3 Pyr neurons is reduced after SAT. (a) Schematic of PV-assigned synaptic puncta analysis for L2/3 Pyr neurons. (b) Representative L2/3 Pyr neuron from ACC animals with PV-assigned (red) and unassigned (green) FAPpost-labeled synapses. Scale bar=20µm. (c) As in (b), but for an L2/3 Pyr neuron after 1 day of SAT (SAT1). (d) Average somatic PV synapse density in L2/3 Pyr neurons from ACC (black) and SAT1 (blue) groups. ACC: 0.065±0.008 /µm^2^ (N=4 mice, n=13 cells); SAT1: 0.038±0.009 /µm^2^ (N=6 mice, n=17 cells). Two-tailed Mann-Whitney U test (U=176, n=13 and 1**7**). (e) Cumulative distribution of somatic PV synapse density in L2/3 Pyr neurons from ACC and SAT1 groups. Kolmogorov-Smirnov (K-S) test. (f-g) As in (d)-(e), but for dendritic PV synapse density. ACC: 0.29±0.04 /µm (N=4 mice, n=13 cells); SAT1: 0.18±0.03 /µm (N=6 mice, n=17 cells). Two-tailed Mann-Whitney U test (U=166, n=13 and 17). (e) K-S test. (h-n) As in (a)-(g), but for L5 Pyr neurons. (k) Somatic density in L5. ACC: 0.041±0.011 /µm^2^ (N=4 mice, n=10 cells); SAT1: 0.041±0.009 /µm^2^ (N=5 mice, n=8 cells). Two-tailed Mann-Whitney U test (U=32, n=10 and 8). (l) K-S test. (m) Dendritic density in L5. ACC: 0.22±0.04 /µm (N=4 mice, n=10 cells); SAT1: 0.19±0.06 /µm (N=5 mice, n=8 cells). Two-tailed Mann-Whitney U test (U=51, n=10 and 8). (n) K-S test. All bar graphs represent mean+SEM. * p<0.05.

In contrast to L2/3, PV synapse density on L5 Pyr neurons did not change (Fig. 2h-n; Soma: ACC 0.041±0.011/µm^2^ vs. SAT1 0.041±0.009/µm^2^, p=0.51; Dendrites: ACC 0.22±0.04/µm vs. SAT1 0.19±0.06/µm, p=0.36). These data suggest that anatomical changes in PV-associated synapses occur selectively in L2/3.

### Training-induced changes in PV synaptic output are female-specific

The anatomical reduction in PV synapses onto neocortical Pyr neurons suggested that PV-mediated inhibitory postsynaptic currents (IPSCs) might also be decreased at the onset of training. To test this, we generated PV-Cre mice that expressed channelrhodopsin (ChR2) in neocortical PV neurons and trained them in the SAT task. A comparison of light-evoked PV-IPSCs revealed a small but significant reduction in L2/3 but not L5 Pyr neurons at the first day of training (Fig. 2d; L2 by cell: ACC 446.0±24.3 pA vs. SAT1 344.4±23.8 pA, p=0.0024; Fig. 2j; L5 by cell: ACC 855.0±57.7 pA vs. SAT1 698.1±59.0 pA, p=0.14). These data are consistent with prior studies indicating that synaptic inhibition onto L2/3 Pyr neurons reduces during the early stages of learning^11,12^.

Recent work suggests that excitability of neocortical PV neurons may be sexually dimorphic, particularly in L5^34–37^, and PV inhibition also differs by sex in hippocampus and ventral subiculum ^38,39^. However, these studies were done in adult animals where the influence of gonadal hormones is high. Sexual dimorphism in PV inhibition in young animals or its learning-related regulation has not been investigated.

To assess this, animals were divided into male and female groups. Prior studies have shown that L5 PV neurons express the estrogen receptor beta (ERβ)^40–42^ and can be regulated by estrogen^42^. Because L5 PV neurons strongly innervate local Pyr neurons^8,43^, we hypothesized that sex-specific PV output modulation may be more pronounced in L5. However, this was not the case. We did not observe a female-specific reduction in PV-IPSCs for L5 Pyr neurons (Fig. 3k-m; ♀ by cell: ACC 990.1±83.9 pA vs. SAT1 722.8±105.6 pA, p=0.089; ♀ by animal: ACC 994.7±96.3 pA vs. SAT1 705.0±145.4 pA, p=0.11; Supplementary Fig. 2; ♂ by cell: ACC 713.1±67.1 pA vs. SAT1 671.4±50.2 pA, p=0.99; ♂ by animal: ACC 732.5±52.1 pA vs. SAT1 683.3±36.5 pA, p=0.54).

**Fig. 3.**
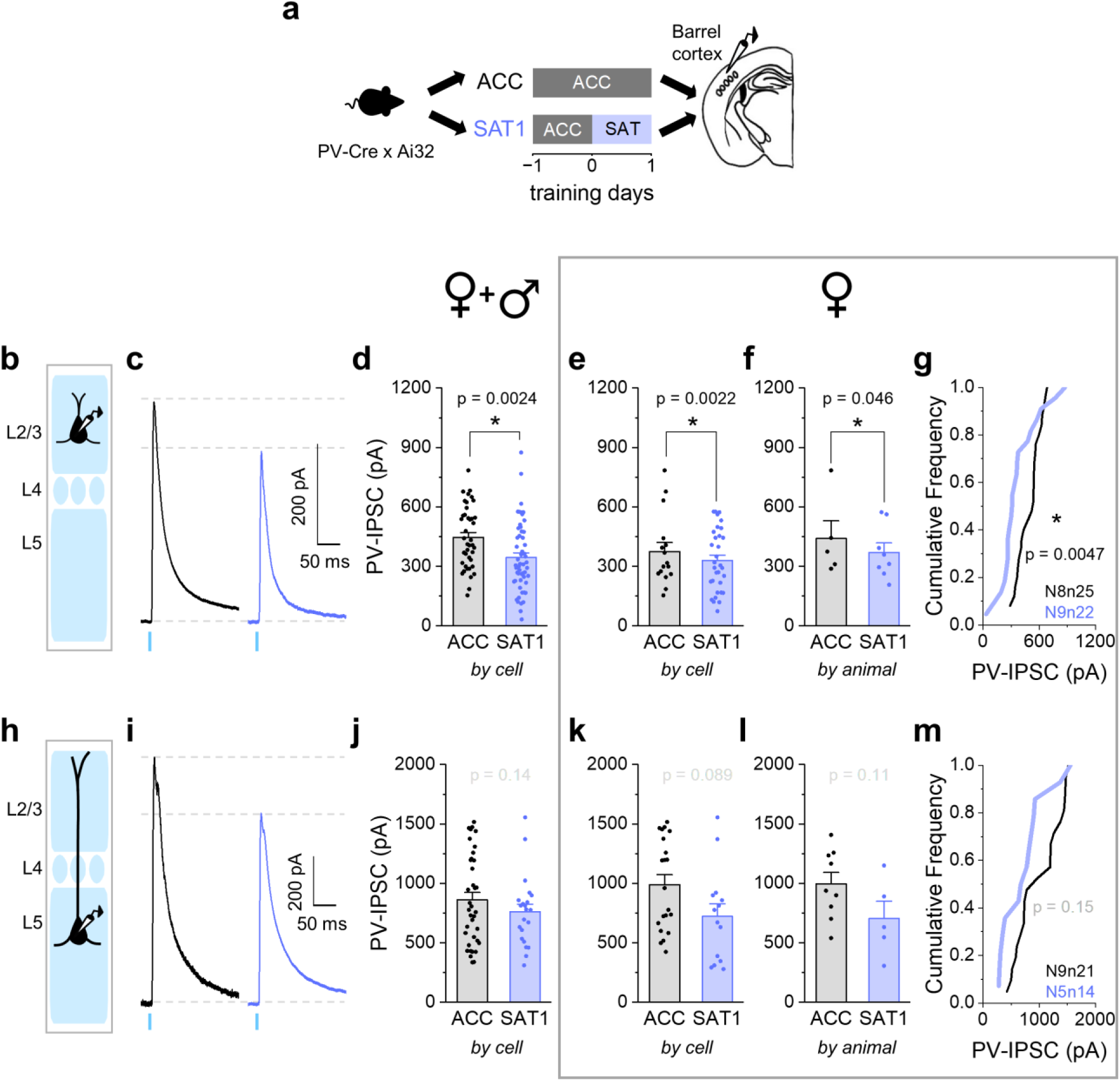
PV-mediated inhibition onto L2/3 Pyr neurons is significantly reduced after one day of training in females. (a) Schematic of training paradigm and tissue preparation for PV-Cre x Ai32 mice. (b) Schematic of whole-cell recordings from L2/3 Pyr neurons. (c) Representative traces of ChR2-evoked PV-IPSCs (10 sweeps average) recorded from L2/3 Pyr neurons after 1-2 days of acclimation (ACC, black) and 1 day of sensory association training (SAT1, blue). (d) Peak PV-IPSC amplitudes in L2/3 Pyr neurons averaged across cells from both male and female mice. ACC: 446.0±24.3 pA (N=13 mice; n=41 cells); SAT1: 344.4±23.8 pA (N=17 mice, n=54 cells). Two-tailed Mann-Whitney U test (U=1506.5, n=41 and 54). (e) Same as in (d), but for cells from females. ACC: 491.3±24.2 pA (N=8 mice, n=25 cells); SAT1: 366.2±42.8 pA (N=9 mice, n=22 cells). Two-tailed Mann-Whitney U test (U=416, n=25 and 22). (f) Same as in (e), but averaged across female mice. ♀ ACC: 515.3±28.7 pA (N=8 mice); ♀ SAT1: 393.5±71.6 pA (N=9 mice). Two-tailed Mann-Whitney U test (U=57, n=8 and 9). (g) Cumulative distribution of PV-IPSC amplitudes in L2/3 Pyr neurons from ACC and SAT1 groups for females. Kolmogorov-Smirnov (K-S) test. (h-m) Same as in (b)-(g), but for L5 Pyr neurons. (j) ACC: 855.0±57.7 pA (N= 15 mice, n=41 cells). SAT1: 698.1±59.0 pA (N=10 mice, n=27 cells). Two-tailed Mann-Whitney U test (U=673, n=41 and 27). (k) ♀ ACC: 990.1±83.9 pA (N=9 mice, n=21 cells); ♀ SAT1: 722.8±105.6 pA (N=5 mice, n=14 cells). Two-tailed Mann-Whitney U test (U=198, n=21 and 14). (l) ♀ ACC: 994.7±96.3 pA (N=9 mice); ♀ SAT1: 705.0±145.4 pA (N=5 mice). Two-tailed Mann-Whitney U test (U=35, n=9 and 5). (m) K-S test. All bar graphs represent mean+SEM. * p<0.05.

Instead, a training-dependent decrease in light-evoked PV-IPSC amplitude in L2/3 Pyr neurons was only apparent in females (Fig. 3e-g; by cell: ACC 491.3±24.2 pA vs. SAT1: 366.2±42.8 pA, p=0.0022; by animal: ACC 515.3±28.7 pA vs. SAT1 393.5±71.6 pA, p=0.046). This was mainly due to larger PV-IPSCs in the control, cage-acclimated females (by cell: ♀ ACC 491.3±24.2 pA vs. ♂ ACC 375.4±44.9 pA, p=0.012) that were reduced by training. Males showed lower PV-IPSCs in both ACC and SAT1 groups (Supplementary Fig. 2; by cell: ACC 375.4±44.9 pA vs. SAT1 329.4±27.7 pA, p=0.52; by animal: ACC 440.1±90.2 pA vs. SAT1 370.7±47.4 pA, p=0.62).

Serum estradiol – a potential source of this sex-specific effect – can start to increase as early as in the 4th postnatal week during development^44,45^, but was not measured in this study. However, we did not observe any correlation of age in the third to fourth postnatal week with L2/3 PV-IPSCs amplitude in our dataset for either sex (Supplementary Fig. 3). Thus, sex-specific regulation of PV inhibition might arise through circuitry established during brain development.

### Sex regulates intrinsic excitability of neocortical PV neurons

The anatomical reduction in PV outputs onto L2/3 Pyr neurons suggested that the change in PV-IPSC amplitude might be explained by the loss of inputs, although we were underpowered to compare sex-dependent changes in these anatomical measurements. Alterations in light-evoked PV firing might also influence recorded PV-IPSCs. To determine whether SAT might drive changes in PV excitability, we compared the intrinsic firing properties of L2/3 and L5 PV neurons across conditions.

For females, SAT was not associated with any change in resting membrane potential of L2/3 PV neurons, although we observed a significant increase in input resistance at SAT1 (Fig. 4d; ACC 141.7±14.4 MΩ vs. SAT1 213.1±19.4 MΩ, p=0.0059). This occurred at the same time as a reduction in rheobase current (Fig. 4e; ACC 190.5±29.1 pA vs. SAT1 112.5±11.4 pA, p=0.023) and a leftward shift in the F:I curve, indicating that L2/3 PV neurons in females were more excitable at the onset of SAT. In contrast to L2/3 PV neurons, intrinsic properties of L5 PV neurons were not altered after SAT (Supplementary Fig. 5). Thus, SAT drives a significant increase in L2/3 PV excitability.

**Fig. 4.**
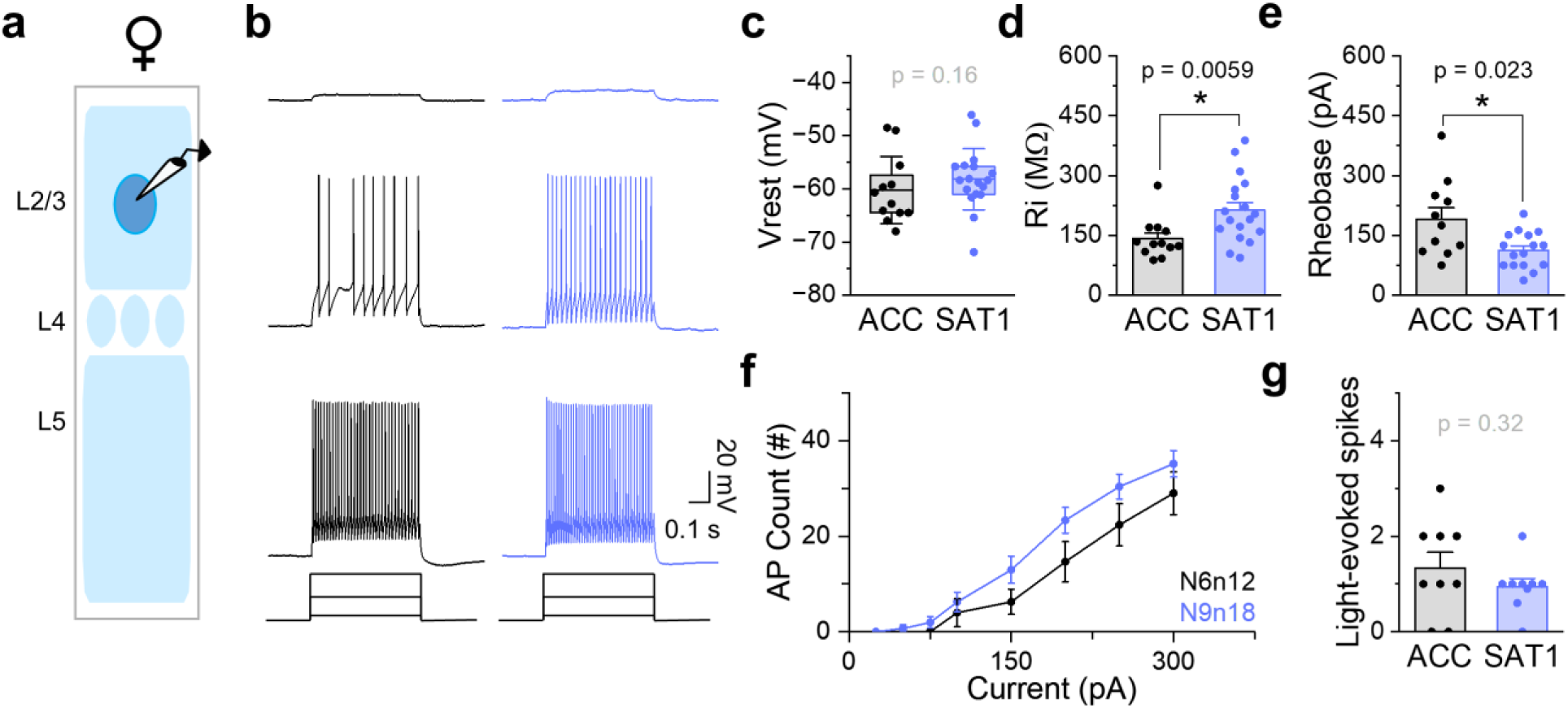
L2/3 PV neuron excitability is increased in females after training. (a) Schematic of the experiment setup. Either eYFP-expressing or ChR2-expressing PV neurons were targeted. (b) Representative firing of L2/3 fast-spiking PV neurons in response to 500 ms current injections (25, 150, 300 pA) after 1-2 days of acclimation (ACC: black) and 1 day of SAT (SAT1: blue). (c) Resting membrane potential (V_rest_) comparison. ACC: −60.2±6.3 mV (N=6 mice, n=12 cells); SAT1: −58.2±5.8 mV (N=9 mice, n=18 cells). The box is the 25th and 75th quartiles, the whiskers are the SD, and the midline is the mean. Two-tailed Mann-Whitney U test (U=74.5, n=12 and 18). (d) Input resistance (Rin) comparison (mean+SEM). ACC: 141.7±14.4 MΩ (N=6 mice, n=12 cells); SAT1: 213.1±19.4 MΩ (N=9 mice, n=18 cells). Two-tailed Mann-Whitney U test (U=44, n=12 and 18). (e) Rheobase comparison (mean+SEM). ACC: 190.5±29.1 pA (N=6 mice, n=11 cells); SAT1: 112.5±11.4 pA (N=9 mice, n=16 cells). Rheobase was recorded from 11 out of 12 cells from ACC mice and 16 out of 18 cells from SAT1 mice. Two-tailed Mann-Whitney U test (U=133.5, n=11 and 16). (f) F-I curve of L2/3 PV neurons in ACC (black) and SAT1 (blue) animals. Mean±SEM. Two-way repeated measures ANOVA F_(1,28)_=2.8, p=0.10. (g) Spiking elicited by brief ChR2 activation (mean+SEM). ACC: 1.3±0.3 (N=5 mice, n=9 cells); SAT1: 0.9±0.5 (N=4 mice, n=9 cells). Two-tailed Mann-Whitney U test (U=52, n=9 and 9). * p<0.05.

If L2/3 PV neurons are more excitable, were they more likely to respond to the light stimulus after SAT? This was not the case, as the probability of spiking and the number of evoked spikes in L2/3 PV neurons were not different between ACC and SAT (Fig. 4g). Overall, the increase in L2/3 PV excitability was not directly related to the magnitude of PV-IPSCs at SAT1, since increased PV excitability or light-evoked PV activation should increase PV-IPSCs. Thus, the female-specific reduction in light-evoked PV inhibition occurs despite increased intrinsic excitability of L2/3 PV neurons.

In males, training increased the rheobase of both L2/3 and L5 PV neurons, indicating reduced intrinsic excitability (Supplementary Fig. 4,6). However, this reduction in excitability was not reflected in PV-IPSC measurements, as light-evoked PV neuron activation remained consistent across conditions (Supplementary Fig. 4,6). These findings indicate that associative learning induces target-specific reduction of PV-mediated inhibition only in females.

### Sex-specific differences in learning behavior

Our original hypothesis was that plasticity of PV inhibition in superficial layers might be an important step in the rewiring of cortical circuits during learning. Since males did not show a training-dependent reduction in PV-IPSCs at the onset of training, we hypothesized that they might be slower to use the stimulus as a cue to increase anticipatory licking during SAT. This was not the case (Fig. 5). Male mice showed a marginally greater increase in anticipatory licking for stimulus trials versus blank trials at the end of the first training day than females, where within-animal comparisons of licking frequency for the last 20% of trials on SAT1 showed a significant difference only for males (Fig. 5c, f; ♀: Stim 4.3±0.4 Hz vs. Blank 3.6±0.4 Hz, p=0.083; ♂: Stim 6.0±0.5 Hz vs. Blank 5.2±0.5 Hz, p=0.016).

**Fig. 5.**
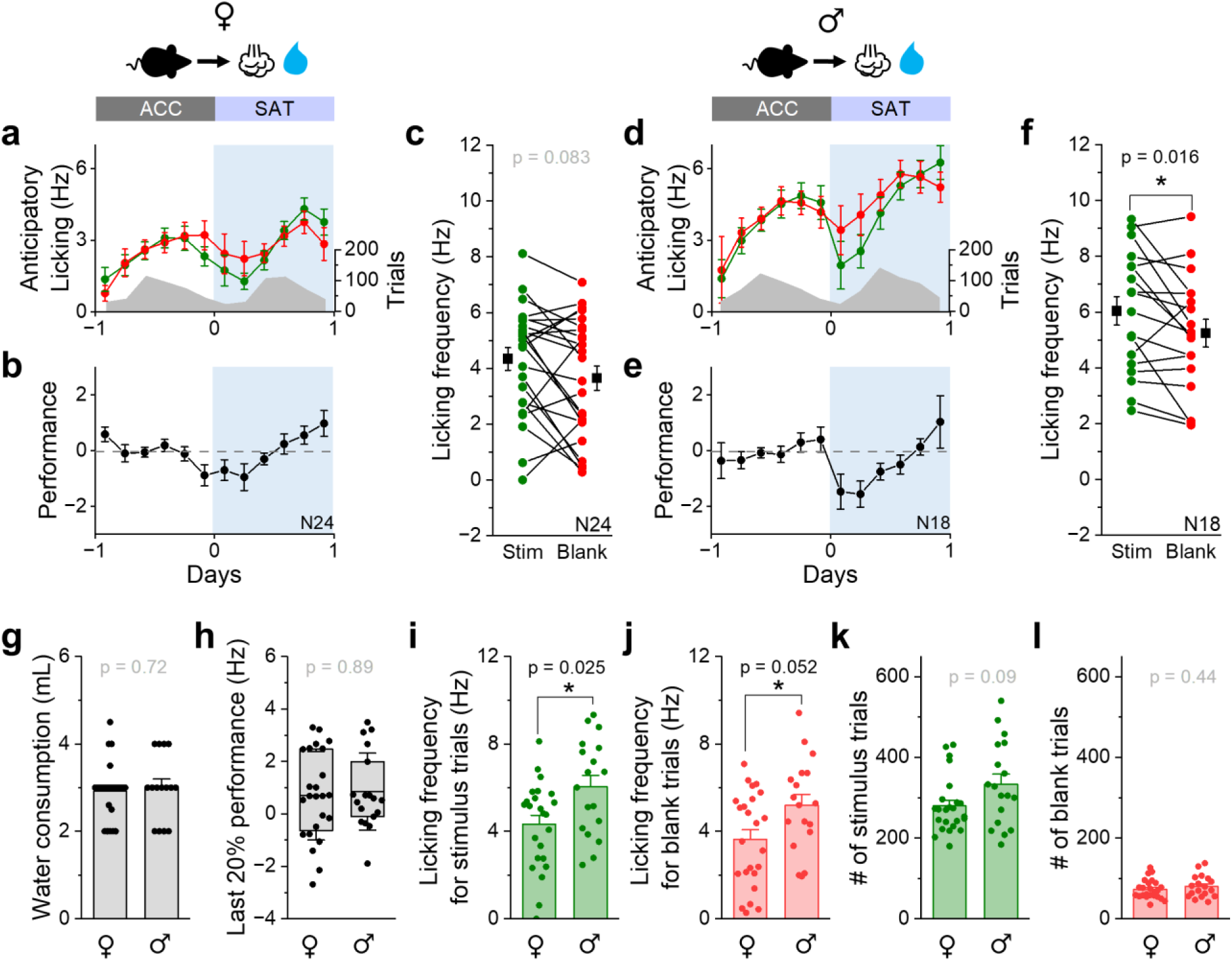
Sex differences in behavior during SAT. (a) Mean anticipatory licking frequency (Hz) for stimulus (green) and blank (red) trials in females after 1 day of SAT (SAT1). The blue shading indicates the training period; the gray shading represents the average trial numbers across time bins. (b) Performance (Lick_Stim_-Lick_Blank_; see Materials and Methods) was calculated for each 4-hr bin and averaged across animals. (c) Mean anticipatory licking frequency (Hz) during the last 20% of stimulus (green) and blank (red) trials for each female animal after SAT1. Black symbols: group mean±SEM. Stim: 4.3±0.4 Hz; Blank: 3.6±0.4 Hz (N=24 mice). Paired-sample, Wilcoxon signed-rank test (Z=1.73, n=24). (d-f) Same as in (a)-(c), but for males. (f) Stim: 6.0±0.5 Hz; Blank: 5.2±0.5 Hz (N=18 mice). *Z*=2.35, n=18. (g) Water consumption during SAT (mean+SEM). ♀: 2.9±0.1 mL (N=23 mice); ♂: 3.0±0.2 mL (N=15 mice). Two-tailed Mann-Whitney U test (U=123.5, n=23 and 15). Data not acquired for one female and three males. (h) Performance during the last 20% of trials. box=25th–75th percentiles; whiskers=SD; midline=mean. ♀: 0.7±1.7 Hz (N=24 mice); ♂: 0.8±1.5 Hz (N=18 mice). U=210, n=24 and 18. (i) Licking frequency for the last 20% stimulus trials (mean+SEM). ♀: 4.3±0.4 Hz (N=24 mice); ♂: 6.0±0.5 Hz (N=18 mice). U=304, n=24 and 18. (j) Licking frequency for the last 20% blank trials (mean+SEM). ♀: 3.6±2.1 Hz (N=24 mice); ♂: 5.2±2.1 Hz (N=18 mice). U=292.5, n=24 and 18. (k) The number of stimulus trials (mean+SEM). ♀: 280±14 trials (N=24 mice); ♂: 334±25 trials (N=18 mice). U=282.5, n = 24 and 18. (l) The number of blank trials (mean+SEM). ♀: 73±5 trials (N=24 mice); ♂: 80±6 trials (N=18 mice). U=247, n=24 and 18. * p<0.05.

To investigate this potential difference, we examined metrics for training behaviors in greater depth. Males showed an overall higher frequency of licking for both stimulus and blank trials at the end of SAT1 (Fig. 5i-j; Stim: ♀ 4.3±0.4 Hz vs. ♂ 6.0±0.5 Hz, p=0.025; Blank: ♀ 3.6±2.1 Hz vs. ♂ 5.2±2.1 Hz, p=0.052), a difference that was highly significant. This increase in licking frequency was not reflected in the overall volume of water consumed during the SAT1 (Fig. 5g; ♀ 2.9±0.1 mL vs. ♂ 3.0±0.2 mL, p=0.72). However, a comparison of performance – defined as the difference in licking frequency between stimulus and blank trials, a metric that can account for animal-to-animal variability in impulsive behavior – was identical between males and females (Fig. 5h; ♀ 0.7±1.7 Hz vs. ♂ 0.8±1.5 Hz, p=0.90). Consistent with a potential increase in impulsive behavior, males initiated a modestly larger number of trials in this freely-moving task, a difference that was not significant (Fig. 5k; ♀ 280±14 trials vs. ♂ 334±25 trials, p=0.09). The correlation between the number of trials and licking frequency was not significant for either male or female mice. Thus, despite a female-specific reduction in L2/3 PV inhibition, learning rates were not enhanced in females. These data are inconsistent with the hypothesis that PV-mediated disinhibition in superficial layers of sensory cortex facilitates association learning in this whisker-dependent task.

### Pseudotraining does not alter PV-inhibition

What properties of SAT drive PV-IPSC plasticity in females during training? Our previous work demonstrated that SST-mediated inhibition in superficial layers is selectively reduced when the stimulus predicts reward^18^. To test whether PV output modulation requires stimulus and reward coupling, we trained a separate group of animals using a pseudotraining (PSE) paradigm, where the stimulus and reward were decoupled but stimulus frequency remained constant (Fig. 6a). PSE animals were compared to age-matched controls that were only acclimated to the training environment.

**Fig. 6.**
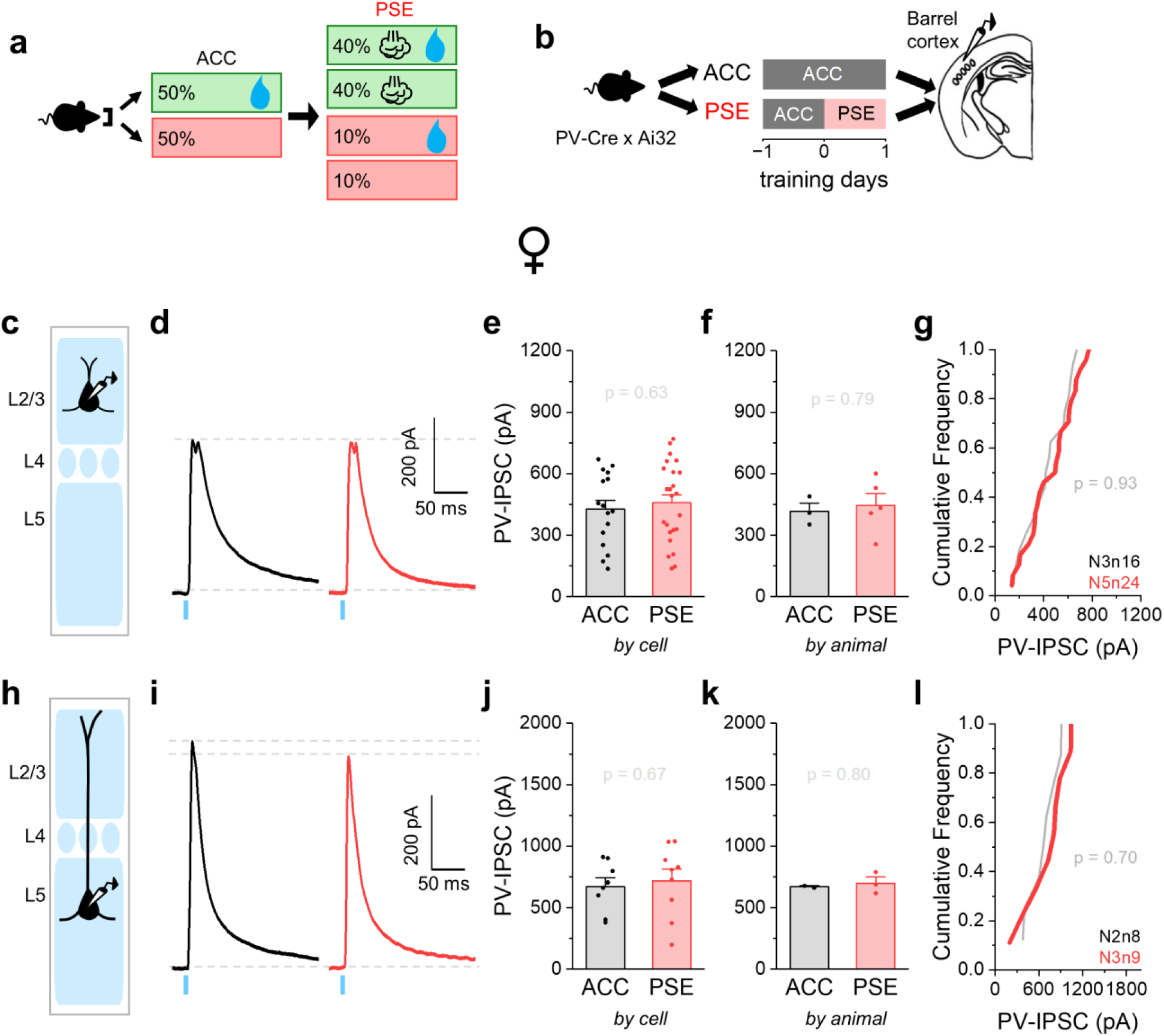
Pseudotraining does not alter PV-mediated inhibition onto Pyr neurons in females. (a) Pseudotraining (PSE) paradigm. (b) Schematic of training paradigm and tissue preparation for PV-Cre x Ai32 mice. (c) Schematic of whole-cell recordings from L2/3 Pyr neurons. (d) Representative traces of ChR2-evoked PV-IPSCs (10 sweeps average) recorded from L2/3 Pyr neurons after 1-2 days of acclimation (ACC, black) and 1 day of PSE (red). (e) Peak PV-IPSC amplitudes in L2/3 Pyr neurons averaged across cells. ACC: 426.7±44.0 pA (N=3 mice, n=16 cells); PSE: 458.7±39.4 pA (N=5 mice, n=24 cells). Two-tailed Mann-Whitney U test (U=174, n=16 and 24). (f) Same as in (e), but averaged across animals. ACC: 417.2±39.7 pA (N=3 mice); PSE: 445.7±58.5 pA (N=5 mice). Two-tailed Mann-Whitney U test (U=6, n=3 and 5). (g) Cumulative distribution of PV-IPSC amplitudes in L2/3 Pyr neurons from ACC and PSE groups. Kolmogorov-Smirnov (K-S) test. (h-l) Same as in (c)-(g), but for L5 Pyr neurons. (j) ACC: 670.4±71.9 pA (N=2 mice, n=8 cells); PSE: 718.3±96.0 pA (N=3 mice, n=9 cells). Two-tailed Mann-Whitney U test (U=31, n=8 and 9). (k) ACC: 670.5±8.2 pA (N=2 mice); PSE: 699.4±49.4 pA (N=3 mice). Two-tailed Mann-Whitney U test (U=2, n=3 and 2). (l) K-S test. All bar graphs represent mean+SEM.

In contrast to SAT, PSE did not drive a change in PV-IPSC amplitude in females for either L2/3 (Fig. 6e-g; by cell: ACC 426.7±44.0 pA vs. PSE 458.7±39.4 pA, p=0.63; by animal: ACC 417.2±39.7 pA vs. PSE 445.7±58.5 pA, p=0.79) or L5 (Fig. 6j-l; by cell: ACC 670.4±71.9 pA vs. PSE 718.3±96.0 pA, p=0.43; by animal: ACC 670.5±8.2 pA vs. PSE 699.4±49.4 pA, p=0.80). Similar to SAT, PSE did not drive PV-IPSC plasticity in male mice (Supplementary Fig. 7). These results suggest that the reduction in PV-mediated inhibition in females requires a consistent stimulus-reward contingency and is not driven by stimulus exposure alone.

### PV-IPSC plasticity is not maintained during longer training periods

Performance continues to increase with longer durations of training (Supplementary Fig. 8). To investigate whether PV-IPSC plasticity in L2/3 or L5 would be enhanced with further training, we examined PV-IPSC amplitude after 5 days of SAT in both male and female mice. To control for potential effects of extended exposure to the homecage training environment, we compared light-evoked PV-IPSCs between control animals that had undergone extended acclimation (ACC6) versus those that had undergone 5 days of SAT (with one day of ACC prior to training). PV-IPSCs at SAT5 were indistinguishable from ACC6 controls in females (Fig. 7), indicating that PV-IPSC plasticity is not sustained throughout the training period, but is transiently evoked at the onset of SAT.

**Fig. 7.**
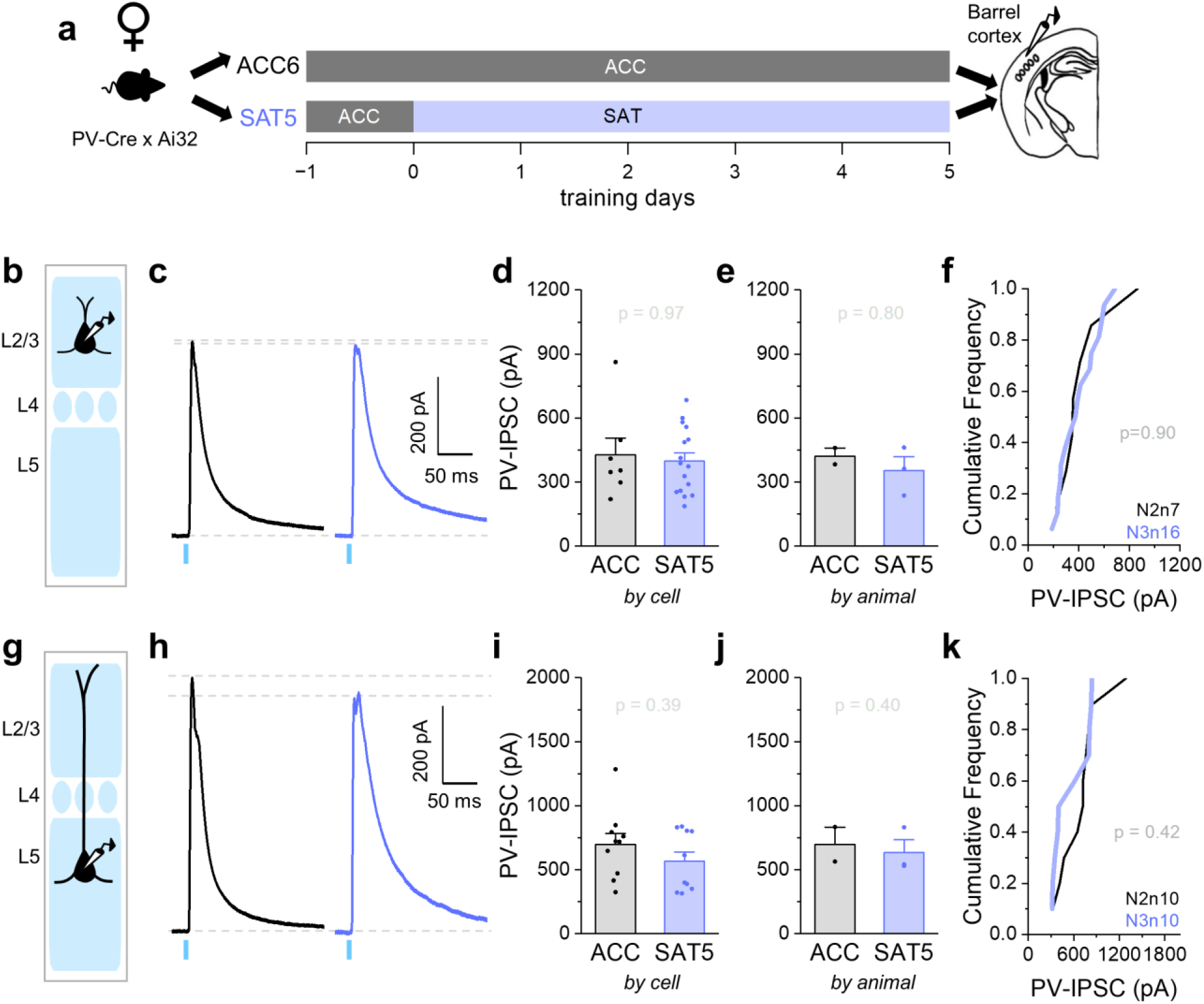
Prolonged training does not alter PV-mediated inhibition onto Pyr neurons in females. (a) Schematic of training paradigm and tissue preparation for PV-Cre x Ai32 mice. (b) Schematic of whole-cell recordings from L2/3 Pyr neurons. (c) Representative traces of ChR2-evoked PV-IPSCs (10 sweeps average) recorded from L2/3 Pyr neurons after 6 days of acclimation (ACC, black) and 5 days of sensory association training (SAT5, blue). (d) Peak PV-IPSC amplitudes in L2/3 Pyr neurons averaged across cells. ACC: 427.0±79.6 pA (N=2 mice, n=7 cells); SAT5: 398.1±38.5 pA (N=3 mice, n=16 cells). Two-tailed Mann-Whitney U test (U=57, n=7 and 16). (e) Same as in (d), but averaged across animals. ACC: 421.5±38.3 pA (N=2 mice); SAT5: 353.7±65.1 pA (N=3 mice). Two-tailed Mann-Whitney U test (U=4, n=2 and 3). (f) Cumulative distribution of PV-IPSC amplitudes in L2/3 Pyr neurons from ACC and SAT5 groups. Kolmogorov-Smirnov (K-S) test. (g-k) Same as in (b)-(f), but for L5 Pyr neurons. (i) ACC: 699.1±85.3 pA (N=2 mice, n=10 cells); SAT5: 566.8±73.5 pA (N=3 mice, n=10 cells). Two-tailed Mann-Whitney U test (U=62, n=10 and 10). (j) ACC: 699.1±134.6 pA (N=2 mice); SAT5: 566.8±635.0 pA (N=3 mice). Two-tailed Mann-Whitney U test (U=5, n=2 and 3). (k) K-S test. All bar graphs represent mean+SEM.

We hypothesized that the threshold for induction of PV inhibitory plasticity might be different between females and males, where males might require a longer period of SAT to initiate PV-IPSC depression. However, we observed no change in PV-IPSCs in either L2/3 or L5 Pyr neurons from male mice at SAT5 (Supplementary Fig. 9). Thus, PV plasticity may be restricted to females in this learning paradigm and this result underscores the importance of sex as a biological variable in understanding how cortical networks respond to stimulus-reward contingencies during learning.

## Discussion

Although the plasticity of inhibitory neurons has long been postulated to have a critical role in learning-related synaptic reorganization^1,11,12,14,15,46^, the input- and target-specific identity of inhibitory plasticity is poorly understood, with few exceptions^18,46–48^. This is critical, since cortical inhibition is mediated by a diverse array of GABAergic neurons with highly selective connectivity^8,49–51^, and inhibitory synaptic plasticity can have very different outcomes depending on the target cell. Because PV neurons are the largest single class of neocortical GABAergic neurons and they densely innervate Pyr neurons to provide rapid and strong feedforward inhibition, decreased PV inhibition might potently regulate learning-related circuit plasticity.

Here, we investigate how association learning can modulate PV-mediated inhibition onto neocortical Pyr neurons. Using patch-clamp recordings of Pyr neurons during light-evoked stimulation of PV neurons, we find that PV-mediated inhibition, particularly onto L2/3 Pyr neurons, is reduced at the onset of training, only in female but not male mice. In addition, SAT was correlated with an increase in the intrinsic excitability of L2/3 PV neurons in females. In males, PV neuron excitability was reduced across the cortical column. Importantly, light-evoked spiking in PV neurons from either sex was not altered by SAT, indicating that changes in PV-IPSCs were not due to altered excitability of PV neurons in either males or females. Both female and male mice learned the association task at similar rates, with small differences in licking frequency between the sexes. These data suggest that sensory association learning can initiate female-specific circuit disinhibition from PV neurons that is most pronounced in the superficial layers of sensory cortex.

Although the anatomical density of PV neurons across the cortex does not differ by sex^52^, sex-specific differences in the intrinsic properties of PV neurons have been observed in higher-order brain areas from adult mice, such as the prefrontal cortex^34–37^. Indeed, PV neurons can express estrogen receptors in both males and females^41,42^. For the juvenile animals used in our study (P23-32), female mice were unlikely to have gone through puberty. Vaginal opening, a precursor to puberty onset, typically occurs after the fourth postnatal week^40,48–51^, and serum estradiol can begin to increase at this time^44,45^. In addition, estradiol can reduce perisomatic inhibition in the female hippocampus^57^. Taken together, it is possible that the early induction of sex hormones in female mice for our study might influence PV-IPSC plasticity. Indeed, in L5 Pyr neurons from control (ACC) mice, PV-IPSCs became smaller with age, a correlation that was significant (Supplementary Fig. 3), suggesting that developmental regulation of sex hormones may be important in regulating PV output onto this layer.

Our study did not investigate serum (or tissue) estradiol levels, so we could not determine whether variations in circulating estrogen might be related to the levels of PV inhibition. Although we were underpowered to investigate sex as a biological variable in our anatomical studies, sexual dimorphism in the number of PV synapses onto some Pyr subtypes has been observed in the ventral subiculum of adult mice^38^. Thus, sex-specific regulation of PV output may not be restricted to sensory neocortex.

Learning-related PV-IPSC plasticity was concentrated in L2/3 Pyr neurons, where there was no significant correlation of PV-IPSCs with age, either for control or trained mice. Importantly, the age range of females across these two groups was identical (26.0 days old; Supplementary Fig. 3), indicating that changes in PV-IPSCs could not be attributed to circuit maturation in trained animals. These data imply that PV plasticity may not be driven by puberty onset in females, although it remains possible that training alters puberty onset, a hypothesis that was not investigated in the present study.

What layer did the specific PV neurons that underwent SAT-dependent plasticity reside? Paired-cell recordings and anatomical reconstructions indicate that L5 PV neurons can project to superficial layers^8,58,59^. Because light-evoked PV-IPSCs can arise from boutons of PV cells across the cortical column, we could not determine the laminar identity of the PV soma that contribute to this SAT-dependent regulation of PV-IPSCs onto L2/3 Pyr neurons.

Acute changes at PV to Pyr synapses have been induced using patterned stimulation in paired recordings from acute brain slices, where PV firing before Pyr neurons is associated with PV-IPSP depression^60^. This spike-timing-dependent plasticity has also been observed in the hippocampus, where PV firing before or at the same time as Pyr neurons also depresses PV output^61^. These data suggest that SAT may engage this pattern of activity to initiate PV-IPSC depression, specifically in females. Interestingly, PSE – where the proportion of stimulus trials was identical to SAT – did not activate this form of synaptic plasticity. Thus, feedforward sensory inputs are not sufficient to drive PV-IPSC changes. These data indicate that the consistency of reinforcement-related signals is critical for the adjustment of PV-IPSC output strength. PV-IPSC synaptic strength may be incrementally adjusted during each trial, where contingent stimulus and rewards depress synaptic output, and stimulus-alone trials reverse this plasticity. Alternatively, predictive sensory information – as established during SAT – may be associated with neuromodulator release that initiates PV-IPSC depression^62^. Future experiments to investigate how dynamic patterns of pre- and postsynaptic activity influence the initiation and stabilization of PV-Pyr synaptic weights may be informative in understanding how SAT and pseudotraining drive different changes in PV-IPSCs.

What are the consequences of reduced PV inhibition in superficial layers? Because PV inhibition is rapidly evoked by sensory stimulation^63,64^, reduced PV inhibition should facilitate recruitment of Pyr neurons, particularly at the onset of sensory stimulation^65^. In line with this, we have previously observed a modest increase in stimulus-evoked Pyr neuron activity at the onset of training^29^. However, this plasticity was not restricted to females, suggesting that changes in PV output may not be required to enhance sensory-evoked responses in L2/3 Pyr neurons.

Reduced PV inhibition can gate excitatory synaptic plasticity^1–3,66^ that can initiate further circuit reorganization associated with learning^67^. Based on this, we hypothesized that layer-specific PV output plasticity might facilitate early potentiation of higher-order thalamocortical inputs onto L5 Pyr neurons occurring during SAT^27,28^. However, since these inputs strengthened similarly in both males and females, we conclude that depression of PV-IPSCs is not required for thalamocortical plasticity in L5. In addition, inhibition onto L5 Pyr neurons from another major class of GABAergic neurons that express SST was not altered by SAT^18^. Thus, thalamocortical plasticity in L5 occurs in the absence of PV- and SST-IPSC depression.

The rapid reduction in PV-IPSCs onto L2/3 Pyr neurons in females occurs simultaneously with the non-sex-specific SST-IPSC depression^18^, and thalamocortical synaptic potentiation is manifested only after several days of training^27^. Our experiments did not determine whether disinhibition is required for this effect. Taken together, these data suggest that disinhibition of L2/3 Pyr neurons may be a critical part of sensory cortex plasticity associated with learning.

Studies have reported that various forms of stress, including social isolation, sleep deprivation, and nerve injury, can induce sex-specific regulation of PV neurons^37,68–72^. It is possible that transferring animals into the training cage differentially alters stress levels according to sex during the acclimation period. Investigating whether PV output onto Pyr neurons is sexually dimorphic outside of this training paradigm may enable a better understanding of sex-dependent circuit properties and their influence on cortical circuit function.

We found that PV-IPSCs are smaller in males during the control, ACC period. Is this associated with enhanced sensory perception in males? In fact, psychophysical studies in humans suggest that females may have greater tactile acuity and higher perceptual sensitivity than males^73–75^. Enhanced tactile sensitivity has also been observed in female mice^76^. On face, it is hard to relate higher perceptual sensitivity in females with enhanced PV inhibition; however, because PV inhibition can regulate spike timing in Pyr neurons as well as control other inhibitory networks^8,49,77^, it is difficult to extrapolate these results to perception. Although the perceptual sensitivity of mice to a whisker stimulus was not directly investigated in this study, our data are consistent with a potential effect of PV-IPSCs on feedforward, stimulus-driven activity. Notably, this putative increase in perceptual sensitivity did not significantly alter learning during SAT, since both males and females showed similar differentiation of anticipatory licking between stimulus and blank trials.

Our data showing sex-specific regulation of PV synaptic output during learning suggests that local computations may be sexually dimorphic and challenges the notion that cortical computations can be generalized from synaptic weights. Computational models of cortical circuits indicate that network stability and output are highly sensitive to PV synapse weights (see for example^78–80^). The 25% difference in mean PV-IPSCs in males vs females in our control group is sufficient to substantially alter network output in some computational models^81,82^. Although this could in theory be balanced by other synaptic changes across the cortical column that were not surveyed in the current study, our findings indicate that cortical networks may be more resilient to changes in synaptic weights than previously thought.

## Materials and Methods

### Animals

All experimental procedures were conducted in accordance with the NIH guidelines and approved by the Institutional Animal Care and Use Committee at Carnegie Mellon University. For functional assessment of PV-to-Pyr synaptic strength, Cre-dependent channelrhodopsin-2 (ChR2; Ai32 Jackson Lab Stock ID 012569^83^) and PV-Cre (Jackson Lab Stock ID 008069^84^) double-transgenic knock-in mice were used (male and female, postnatal day (P)25-29). For a subset of experiments assessing PV neuron excitability, we used PV-tdTomato mice (Jackson Lab Stock ID 027395) and double-transgenic PV-Cre × Ai3 (Cre-dependent YFP; Jackson Lab Stock ID 007903^85^) mice to target PV neurons. PV-Cre, Ai32, Ai3 and PV-tdTomato mice were obtained from Jackson Laboratory.

For anatomical experiments, barrel cortex from double-transgenic PV-Cre x Ai3 was stereotaxically injected with FAPpost (0.1µL), a neuroligin1-based rAAV construct that mediates far-red fluorescent signal at postsynaptic sites^31^. Virus was introduced through a small craniotomy (from bregma: x=-3, y=-0.9, z=-0.5 mm) using a Nanoject II (Drummond Scientific Company; Broomall, PA) in isoflurane-anesthetized mice at P15-17. Six to 8 days later, animals underwent whisker-stimulation reward association training. We confirm that the present study was performed in accordance with ARRIVE guidelines.

### Automated sensory association training

We used an automated, high-throughput experimental paradigm for multiwhisker stimulus-reward training for sensory learning as described previously^27^. Briefly, animals were housed in modified homecages equipped with an SAT chamber in which initiating nosepokes at the waterport caused an infrared beam break that triggered trial onset with a random variable delay (0.2-0.8 s) preceding the whisker stimulus. Animals received all their water from this waterport and were not otherwise water-restricted. During SAT, 80% of stimulus trials began with the administration of a gentle, downward-projecting airpuff directed against right-side whiskers (4-6 PSI, 0.5 s duration). One second after trial onset, a water reward (∼8-25 µL) was delivered to the lickport. For the remaining 20% of blank trials, nosepokes triggered an approximately 2-3 s timeout (depending on random delay duration; Fig. 1). During pseudotraining, airpuff stimulation was administered in 80% of stimulus trials, and water was delivered for half of those trials. For the remaining 20% of blank trials, water was delivered for half of those trials. Thus, in SAT experiments, airpuff was predictive of water reward, and in pseudotraining experiments, sensory stimulation was uncoupled from water reward. For SAT and pseudotraining experiments, cage-matched controls were used. Performance was calculated as the difference in anticipatory lick rates (0.3 sec prior to water delivery) for stimulus trials vs. blank trials (Lick_Water_-Lick_Blank_). Blank trial anticipatory licking was assessed in the 0.3 s preceding water delivery. This occurred within 0.9-1.5 s following the nosepoke, where variation was due to the random delay between nosepoke and trial onset/water delivery. Mean anticipatory lick rates for each animal were calculated in 4-hour bins, from SAT-chamber acclimation 1 day before experiment onset through to the end of the experiment. Animals were excluded from the study if they engaged in <50 trials or consumed <1mL of water in the 1 day preceding electrophysiological or anatomical analysis. For reliable estimates of performance, we required that a minimum of 10 total trials (stimulus and blank trials) within a 4-hour window had to be completed to include data from an individual animal.

### Electrophysiology

At midday (11am-2pm), following SAT (SAT1 or SAT5) or housing in training cages without airpuff exposure (ACC1-2 or ACC6), mice (P25-29) were briefly anesthetized with isoflurane before decapitation. Angled-coronal slices (45° rostro-lateral; 350 µm thick) designed to preserve columnar connections in barrel cortex were prepared in ice-cold artificial cerebrospinal fluid (ACSF) composed of (in mM): 119 NaCl, 2.5 KCl, 1 NaH_2_PO_4_, 26.2 NaHCO_3_, 11 glucose, 1.3 MgSO_4_, and 2.5 CaCl_2_ equilibrated with 95%O_2_/5%CO_2_. Slices were allowed to recover at room temperature in ACSF for one hour in the dark before targeted whole-cell patch-clamp recordings were performed using an Olympus light microscope (BX51WI) and borosilicate glass electrodes (4-8 MΩ resistance) filled with internal solution composed of (in mM): 125 potassium gluconate, 10 HEPES, 2 KCl, 0.5 EGTA, 4 Mg-ATP, 0.3 Na-GTP, and trace amounts of AlexaFluor 594 or AlexaFluor 568 (pH 7.25-7.30, 290 mOsm). Because of the need to verify cell type identity using the action potential waveform, we used a K-gluconate-based internal solution to enable spiking. Electrophysiological data were acquired using a MultiClamp 700B amplifier, digitized with a National Instruments acquisition interface, and collected using MultiClamp and IgorPro6.0 software with 3kHz filtering and 10 kHz digitization. L2/3 and L5 Pyr neurons were targeted based on Pyr morphology, using the pial surface and dense PV-Ai32 fluorescence in L4 barrels for laminar orientation.

Following whole-cell break-in, presumptive Pyr cell identity was confirmed based on hyperpolarized resting membrane potential (approximately −70mV in L2/3 and −60mV in L5), input resistance (approximately 100-200 MΩ; < 400MΩ cut-off), and regular-spiking (RS) action potential waveforms recorded in responses to progressive depolarizing current injection steps recorded in current-clamp mode (50-400 pA, Δ50 pA steps, 0.5s duration). L5 Pyr neurons were typically in the top to the middle portion of L5 (L5a) and had either an RS or intrinsically bursting (IB) phenotype with current injection. Only cells with a stable baseline holding potential, resting membrane potential <-50mV, and access resistance <40MΩ were analyzed^86^. Similar to previously published studies^23,49^, we isolated PV-mediated IPSCs using transgenic mice expressing channelrhodopsin in all presynaptic PV neurons using K-gluconate internal and a depolarized holding potential to evoke outward currents. We selected a holding potential (−50 mV) from the middle of their range (−45 to −55 mV). Blue light stimulation was used to evoke PV-IPSCs (480nm, ∼0.48 mW, 5 ms pulse). Initially, PV-IPSCs were tested at a range of LED light intensities (0.02 – 2.7 mW). We selected 0.48 mW because it evoked IPSC amplitudes below 1 nA (minimizing voltage-clamp errors) and reliably elicited IPSCs in all trials. Direct recording from PV neurons indicated that this intensity was sufficient to drive at least one spike in >75% of cells, a property that was not affected by SAT. Consistent with a chloride-mediated current, the reversal potential for optically-evoked currents was experimentally determined to be −78±4 mV (uncorrected for junction potential). Five minutes after the break-in, Pyr cells were voltage-clamped (VC) at −50 mV, PV-mediated IPSCs were collected using light stimulation, and peak amplitude was assessed using the maximum peak of the evoked response. Because optogenetic activation of PV outputs does not synchronously activate all presynaptic neurons, IPSCs could show multiple peaks in some cells. Recordings were performed blinded to the experimental condition for a subset of experiments.

To assess L2/3 PV neuron excitability in current clamp recordings, PV neurons were targeted in either PV-Cre x Ai32, PV-Cre x Ai3, or PV-tdTom transgenic mouse tissue^87^. PV neuron identity was verified by reporter fluorescence, fast-spiking phenotype in response to direct depolarizing current injection, and/or the presence of excitatory photocurrents in response to blue light stimulation. Rheobase and evoked spiking were assessed using progressive depolarizing current injection steps recorded in current-clamp mode (25-300 pA, Δ25-50 pA steps, 0.5s duration). Only PV cells with a stable baseline holding potential and resting membrane potential <-45mV were included in the dataset for analysis. An LED power of 0.48 mW (5 ms, the same light power used to evoke PV-IPSCs) was used to quantify the number of light-evoked spikes.

### Anatomy

At midday following 1 day of SAT, animals were anesthetized with isoflurane and transcardially perfused using 20 mL PBS (pH 7.4) followed by 20 mL 4% paraformaldehyde (PFA) in PBS (PFA; pH 7.4). Brains were removed and postfixed overnight at 4°C in 4% PFA before transfer into 30% sucrose/PBS. After osmotic equilibration, 45 µm-thick brain sections were collected using a freezing microtome. Free-floating brain sections containing dTom-expressing cells in the barrel cortex were washed with PBS before 30-minute room temperature incubation with MG-Tcarb dye (300nM in PBS) for activation of the far-red fluorescence of the FAP^88^.

Pyr neurons were identified by their pyramid-shaped cell body, a narrow axon descending from the base of their soma, a prominent apical dendrite, and laterally projecting, spiny basal dendrites. Confocal image stacks centered around a well-isolated, FAPpost-expressing Pyr soma were collected with an LSM 880 AxioObserver Microscope (Zeiss) using a 63x oil-immersion objective lens (Plan-Apochromat, 1/40 Oil DIC M27) with the zoom factor set to 1 and the pinhole set at 1.0 Airy disk unit for each fluorescence channel. Optimal laser intensities for each channel were independently set to avoid pixel saturation for each cell. Fluorescence acquisition settings were as follows: YFP (excitation λ514, detection λ517–535), dTom (excitation λ561, detection λ561–597), and MG/FAP (excitation λ633, detection λ641–695). Maximum image size was 1024×1024 pixels, to collect 135 x 135 x ≤ 45µm images, with corresponding 0.13 x 0.13 x 0.3µm voxel dimensions.

Synapse distribution analysis was carried out using previously published methods for the FAPpost synaptic marker^31^. In brief, Carl Zeiss image files were imported into Imaris (v8.4 with FilamentTracer; Bitplane; Zürich, Switzerland) and the dTom cell fill was used to create a 3D Pyr neuron rendering using Imaris macros to create a combination of “surface” and “filament” objects. FAPpost puncta were then reconstructed as “surfaces” using an estimated 0.5µm diameter, 4-voxel minimum, and a spit-touching object setting using the same 0.5µm diameter. FAPpost “surfaces” were digitally assigned to a given neuron if their edges lay within 0.5µm of the soma surface (inner and outer edge), or ≤1µm from a dendrite. Puncta “surfaces” were converted into puncta “spots” (created using automatic intensity-maxima background-subtraction thresholds with an estimated 0.5µm diameter) using “surface” object centroids. Presynaptic neurite reconstructions were created using automatic background-subtraction thresholding of presynaptic PV-YFP fluorescence using an estimated diameter of 0.6µm, split-touching object diameter threshold of 1µm (applied with automatic “quality” filter setting), and a 1µm2 minimum surface area. To digitally correct for z-axis-related signal drop-off, neurite reconstructions using automatic settings were generated separately for every 10µm of z-depth, resulting in similar density and size profiles for both superficial and deep presynaptic neurite reconstructions. Finally, FAPpost puncta “spots” were assigned as PV+ using a distance threshold of 0.15µm from the spot centroid to presynaptic neurite edge (PV synapse).

Since discrete classes of PV neurons may differentially target Pyr neuron compartments^89–92^, compartment-specific methods for assessing PV synapses were used to serve as a guide for evaluating whether a specific population of presynaptic PV neurons might be differentially affected by SAT. During preliminary analysis, PV synapse density across Pyr dendrites was assessed separately for apical and basal dendrite segments (across branch orders), soma, and axon compartments by taking the total number of PV-assigned synapses for each compartment and dividing it by the total length (for dendrites and axon) or surface area (for soma). Since a similar decrease in PV-assigned synapse density was observed across all dendritic compartments (low and higher order apical and basal dendrites), all dendritic compartments were pooled in the final analysis, and reported densities were calculated using the total number of spots (total FAPpost and PV synapses) divided by the total length of the dendrite analyzed.

### Statistics

Mean anticipatory lick-rate and performance (±SEM) for each 4-hour time bin were calculated to evaluate average group behavior. Changes in water and blank trial mean anticipatory licking for the last 20% of trials performed in a given training day were assessed using the paried-sample, Wilcoxon signed-rank test. PV-IPSC magnitudes, membrane potential, input resistance, rheobase current, optically-evoked spike counts, as well as PV neurite and synapse densities (per µm for dendrite and per µm^2^ for soma) were assessed for statistical significance using the Mann-Whitney U test and the Kolmogorov-Smirnov (K-S) test. Sex differences in water consumption, performance, licking frequency for stimulus and blank trials, and the number of stimulus and blank trials were assessed across sexes using the Mann-Whitney U test. Independent control groups were used for SAT1, SAT5, and pseudotraining conditions, and comparisons between control and experimental groups were made within layer (L2/3 or L5). Unless otherwise noted, IPSC amplitudes, excitability measures, PV neurite, and PV synapse densities averaged by cell are reported in the text and represented in graphs as mean±SEM (with individual cell values overlaid). The effect of the current injection step and experimental conditions on firing frequency responses was assessed using two-way ANOVA. All statistical tests and IPSC peak analysis were performed using OriginPro 2024b software (Northampton, MA). Statistical significance, *p* < 0.05.

## Data Availability

The datasets generated during the current study are available from the corresponding author on reasonable request.

## Acknowledgements

Special thanks to Joanne Steinmiller and Rachel Bouchard for expert management of transgenic mice, Sarah Bernhard for SAT cage design and technical support, Ajit Ray and Alex Hsu for custom MatLab scripts for behavioral analysis, Marcel Bruchez for providing reagents for FAPpost detection, and members of the Barth Lab for helpful comments on the manuscript.

## Author contributions

EP, DAK, SEM, and JAC acquired and analyzed electrophysiological data, and EP and DAK acquired and analyzed anatomical data. Experimental design and data interpretation were performed by EP, DAK, and ALB. All authors contributed to writing the manuscript.

## Additional information

The authors declare no competing interests.

