## Supplementary Figures for "Sexually dimorphic plasticity of PV inhibition in sensory neocortex during learning"

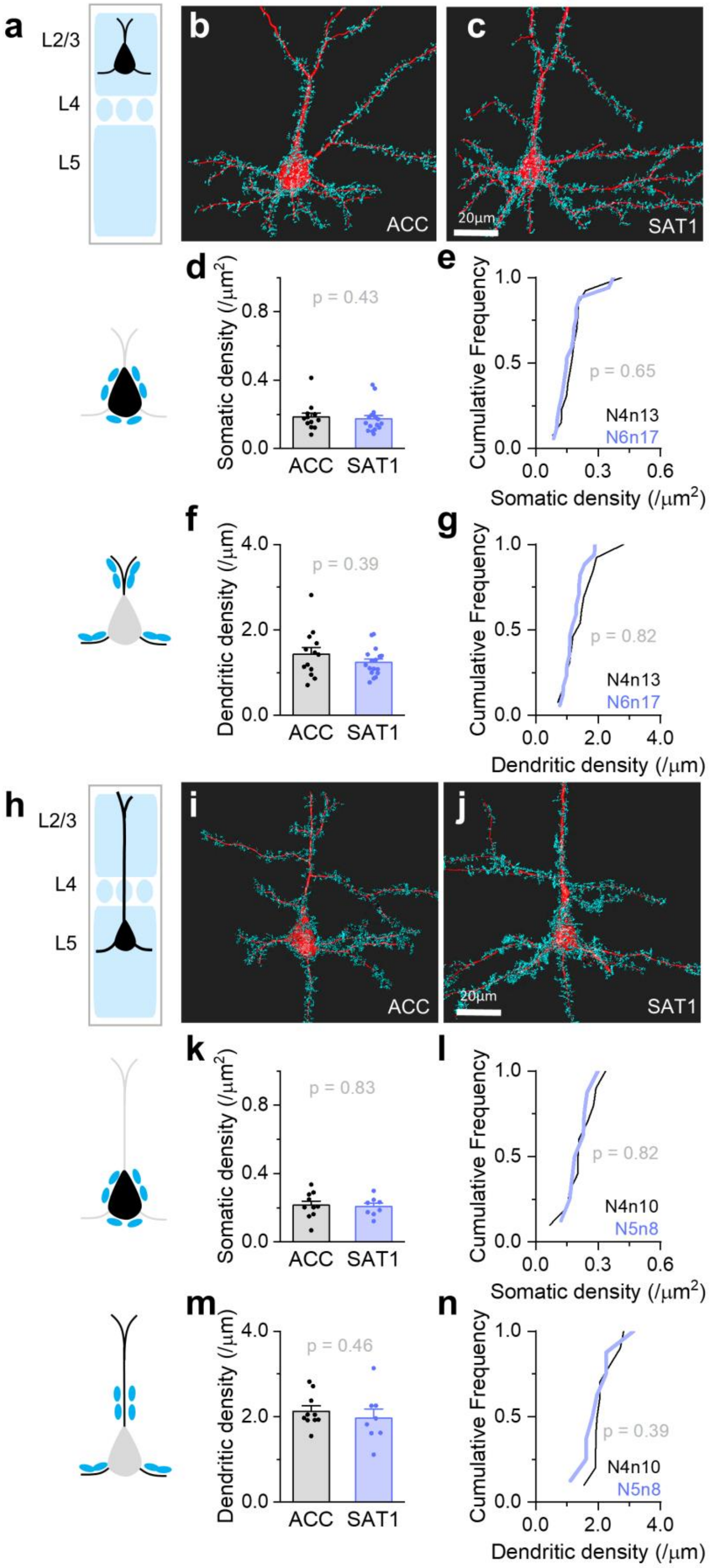


**Supplementary Fig. 1. SAT does not alter presynaptic PV neurite density associated with L2/3 and L5 Pyr neurons.**

(a) Schematic of L2/3 anatomical analysis. (b) Representative L2/3 Pyr neuron (red) with nearby PV neurites (cyan) from ACC animals. Scale bar = 20µm. (c) As in (b), but for an L2/3 Pyr neuron after 1 day of SAT (SAT1). (d) Average Somatic PV neurite density in L2/3 Pyr neurons from ACC (black) and SAT1 (blue) groups. ACC: 0.19 ± 0.02 /µm$^2$ (N = 4 mice, n = 13 cells); SAT1: 0.17 ± 0.02 /µm$^2$ (N = 6 mice, n = 17 cells). Two-tailed Mann-Whitney U test (U = 130, n = 13 and 17). (e) Cumulative distribution of somatic PV neurite density in L2/3 Pyr neurons from ACC and SAT1 groups. Kolmogorov-Smirnov (K-S) test. (f-g) As in (d)-(e), but for dendritic PV neurite density. ACC: 1.4 ± 0.2 /µm (N = 4 mice, n = 13 cells); SAT1: 1.2 ± 0.1 /µm (N = 6 mice, n = 17 cells). Two-tailed Mann-Whitney U test (U = 132, n = 13 and 17). (e) K-S test. (h-n) As in (a)-(g), but for L5 Pyr neurons. (k) Somatic PV neurite density in L5. ACC: 0.21 ± 0.02 /µm$^2$ (N = 4 mice, n = 10 cells); SAT1: 0.21 ± 0.02 /µm$^2$ (N = 5 mice, n = 8 cells). Two-tailed Mann-Whitney U test (U = 43, n = 10 and 8). (l) K-S test. (m) Dendritic PV neurite density in L5. ACC: 2.1 ± 0.1 /µm (N = 4 mice, n = 10 cells); SAT1: 2.0 ± 0.2 /µm (N = 5 mice, n = 8 cells). Two-tailed Mann-Whitney U test (U = 49, n = 10 and 8). (n) K-S test. All bar graphs represent mean + SEM.

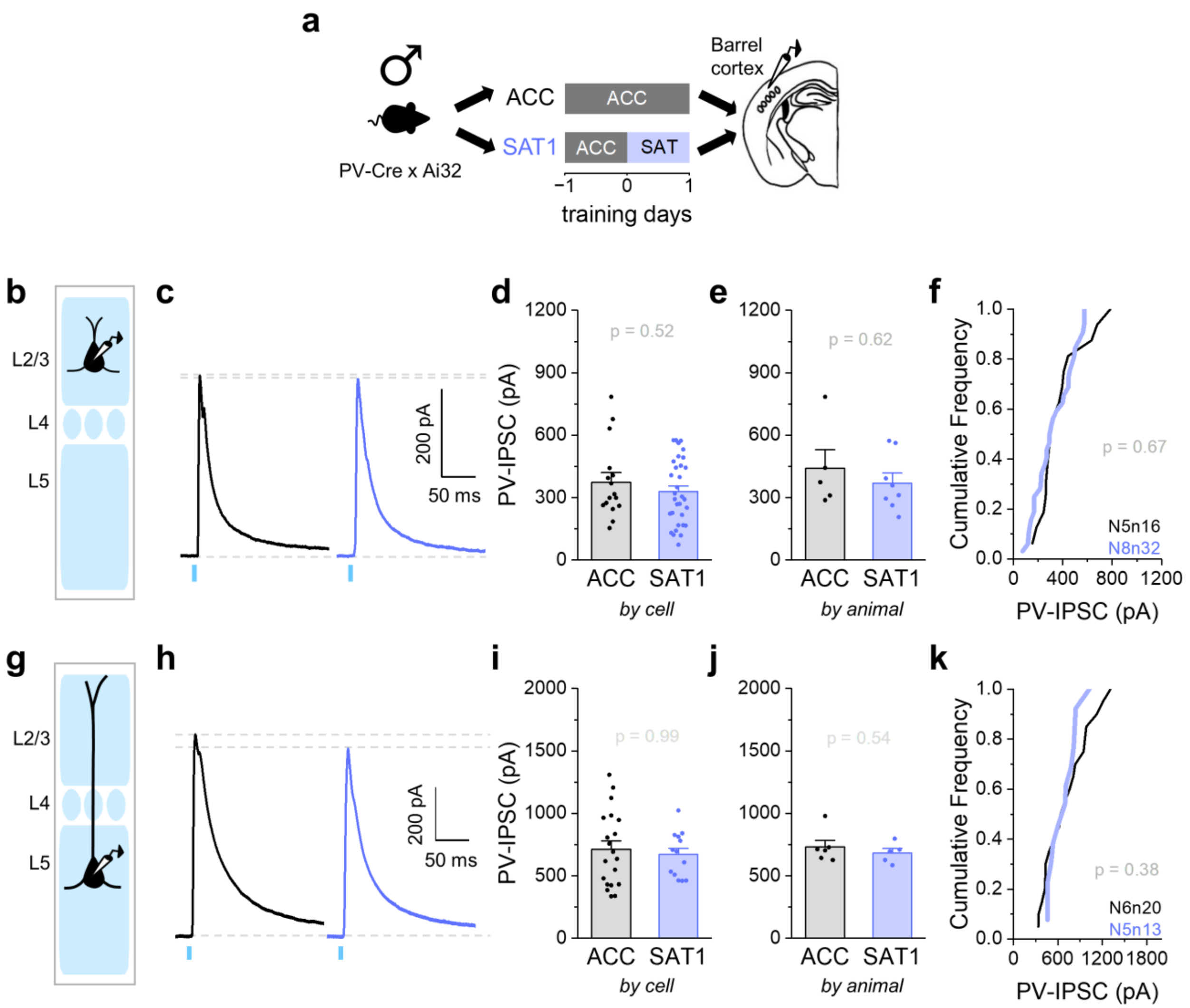


**Supplementary Fig. 2. PV-mediated inhibition onto L2/3 Pyr neurons is not altered after one day of training in males.**

(a) Schematic of training paradigm and tissue preparation for PV-Cre x Ai32 mice. (b) Schematic of whole-cell recordings from L2/3 Pyr neurons. (c) Representative traces of ChR2-evoked PV-IPSCs (10 sweeps average) recorded from L2/3 Pyr neurons after 1-2 days of acclimation (ACC, black) and 1 day of sensory association training (SAT1, blue). (d) Peak PV-IPSC amplitudes in L2/3 Pyr neurons averaged across cells. ACC: 375.4 ± 44.9 pA (N = 5 mice, n = 16 cells); SAT1: 329.4 ± 27.7 pA (N = 8 mice, n = 32 cells). Two-tailed Mann-Whitney U test (U = 286, n = 16 and 32). (e) Same as in (d), but averaged across animals. ACC: 440.1 ± 90.2 pA (N = 5 mice); SAT1: 370.7 ± 47.4 pA (N = 8 mice). Two-tailed Mann-Whitney U test (U = 24, n = 5 and 8). (f) Cumulative distribution of PV-IPSC amplitudes in L2/3 Pyr neurons from ACC and SAT1 groups. Kolmogorov-Smirnov (K-S) test. (g-k) Same as in (b)-(f), but for L5 Pyr neurons. (i) ACC: 713.1 ± 67.1 pA (N = 6 mice, n = 20 cells); SAT1: 671.4 ± 50.2 pA (N = 5 mice, n = 13 cells). Two-tailed Mann-Whitney U test (U = 129, n = 20 and 13). (j) ACC: 732.5 ± 52.1 pA (N = 6 mice); SAT1: 683.3 ± 36.5 pA (N = 5 mice). Two-tailed Mann-Whitney U test (U = 19, n = 6 and 5). (k) K-S test. All bar graphs represent mean + SEM. * p<0.05.

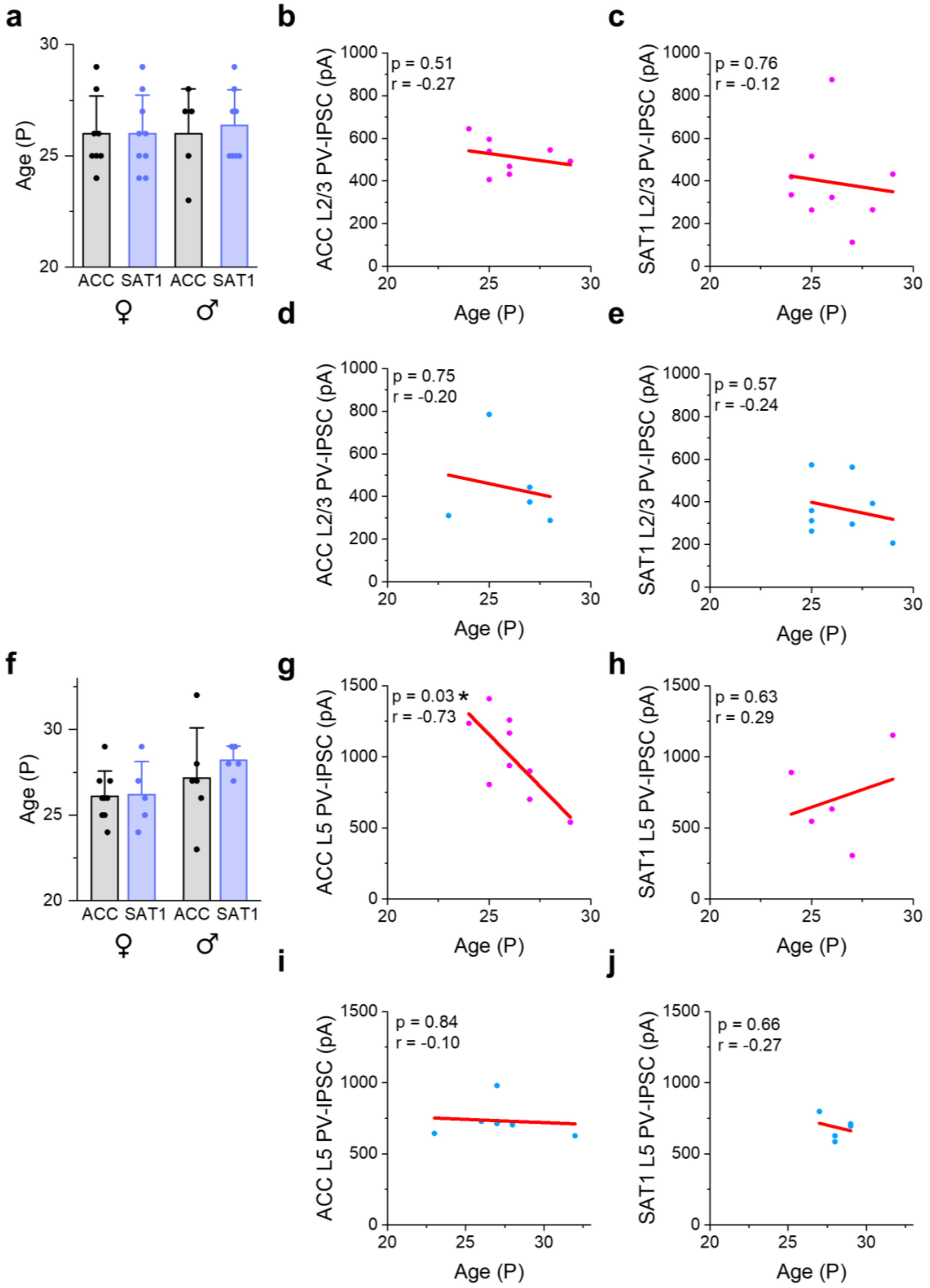


**Supplementary Fig. 3. L2/3 PV-IPSCs are not correlated with age in either sex.**
(a) Average age of animals used for L2/3 PV-IPSC recordings shown in Fig. 3 and Supplementary Fig. 2. ♀ ACC: P26.0 ± 1.7 (N = 8 mice); ♀ SAT1: P26.0 ± 1.7 (N = 9 mice); ♂ ACC: P26.0 ± 2.0 (N = 5 mice); ♂ SAT1: P26.4 ± 1.6 (N = 8 mice). (b) Correlation between L2/3 PV-IPSC amplitude and age in female mice from the acclimation (ACC) group. (c) Same as in (b), but for female mice in the SAT1 group. (d–e) Same

as in (b–c), but for male mice. (f-j) Same as in (a)-(e), but for animals used for L5 PV-IPSC recordings shown in Fig. 3 and Supplementary Fig. 2. (f) ♀ ACC: P26.1 ± 1.5 (N = 9 mice); ♀ SAT1: P26.2 ± 1.9 (N = 5 mice); ♂ ACC: P27.2 ± 2.9 (N = 6 mice); ♂ SAT1: P28.2 ± 0.8 (N = 5 mice). Bar graphs represent mean + SD. Pearson's correlation. * $p < 0.05$.

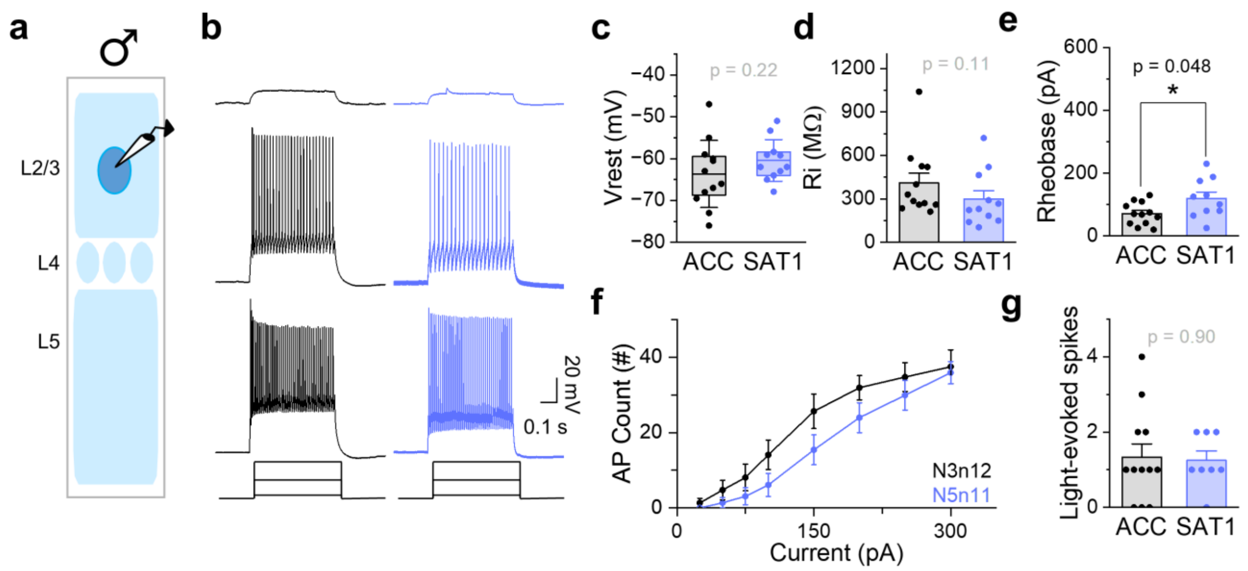


**Supplementary Fig. 4. Excitability of L2/3 PV neurons in males is unaltered after training.**
(a) Schematic of the experiment setup. Either eYFP-expressing or ChR2-expressing PV neurons were targeted. (b) Representative firing of L2/3 fast-spiking PV neurons in response to 500 ms current injections (25, 150, 300 pA) after 1-2 days of acclimation (ACC: black) and 1 day of SAT (SAT1: blue). (c) Resting membrane potential (Vrest) comparison. ACC: -63.7 ± 8.0 mV (N = 3 mice, n = 12 cells); SAT1: -60.4 ± 5.0 mV (N = 5 mice, n = 11 cells). The box is the 25th and 75th quartiles, the whiskers are the SD, and the midline is the mean. Two-tailed Mann-Whitney U test (U = 45.5, n = 12 and 11). (d) Input resistance (Rin) comparison (mean + SEM). ACC: 409.8 ± 67.9 MΩ (N = 3 mice, n = 12 cells); SAT1: 299.1 ± 57.5 MΩ (N = 5 mice, n = 11 cells). Two-tailed Mann-Whitney U test (U = 92.5, n = 12 and 11). (e) Rheobase comparison (mean + SEM). ACC: 70.8 ± 10.4 pA (N = 3 mice, n = 12 cells); SAT1: 119.9 ± 19.9 pA (N = 4 mice, n = 10 cells). Rheobase was recorded from 10 out of 11 cells from SAT1 mice. Two-tailed Mann-Whitney U test (U = 30, n = 12 and 10). (f) F-I curve of L2/3 PV neurons in ACC (black) and SAT1 (blue) animals. Mean ± SEM. Two-way repeated measures ANOVA $F_{(1,21)} = 2.3$, $p = 0.15$. (g) Spiking elicited by brief ChR2 activation (mean + SEM). ACC: 1.3 ± 0.4 (N = 3 mice, n = 12 cells); SAT1: 1.3 ± 0.3 (N = 3 mice, n = 8 cells). Two-tailed Mann-Whitney U test (U = 45.5, n = 12 and 8). * $p<0.05$.

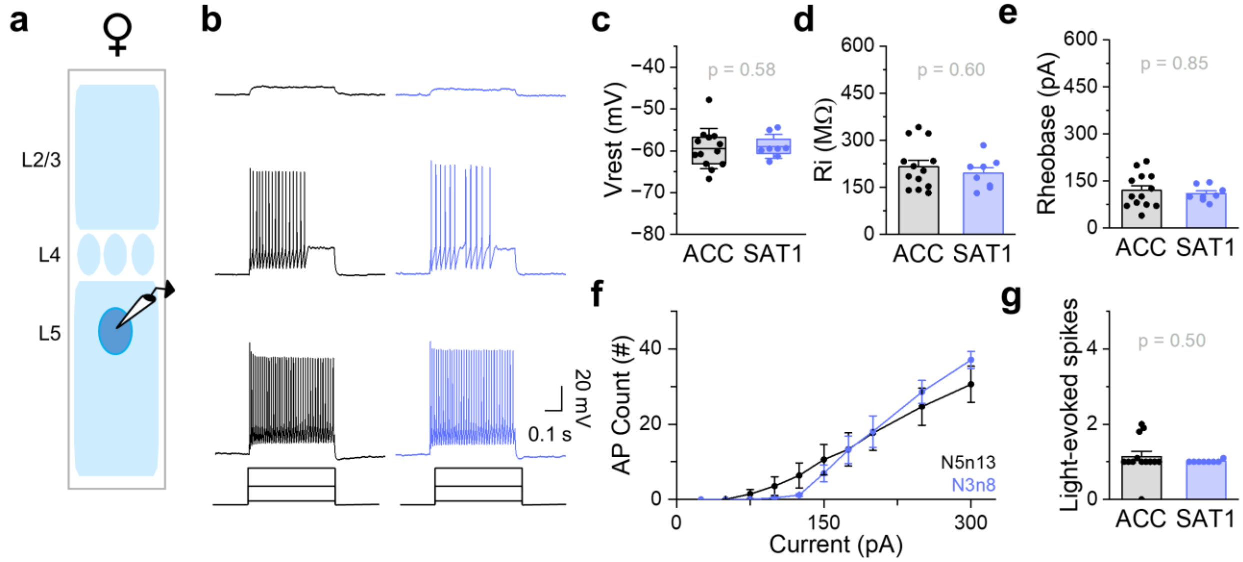


**Supplementary Fig. 5. Intrinsic properties of L5 PV neurons in females are unaltered after training.** (a) Schematic of the experiment setup. ChR2-expressing L5 PV neurons were targeted. (b) Representative firing of L5 fast-spiking PV neurons in response to 500 ms current injections (25, 150, 300 pA) after 1-2 days of acclimation (ACC: black) and 1 day of SAT (SAT1: blue). (c) Resting membrane potential (Vrest) comparison. ACC: -59.5 ± 4.8 mV (N = 5 mice, n = 13 cells); SAT1: -59.0 ± 2.9 mV (N = 3 mice, n = 8 cells). The box is the 25th and 75th quartiles, the whiskers are the SD, and the midline is the mean. Two-tailed Mann-Whitney U test (U = 44, n = 13 and 8). (d) Input resistance (Rin) comparison (mean + SEM). ACC: 215.7 ± 20.2 MΩ (N = 5 mice, n = 13 cells); SAT1: 195.3 ± 17.8 MΩ (N = 3 mice, n = 8 cells). Two-tailed Mann-Whitney U test (U = 60, n = 13 and 8). (e) Rheobase comparison (mean + SEM). ACC: 120.0 ± 15.1 pA (N = 5 mice, n = 13 cells); SAT1: 109.9 ± 8.6 pA (N = 3 mice, n = 8 cells). Two-tailed Mann-Whitney U test (U = 55, n = 13 and 8). (f) F-I curve of L5 PV neurons in ACC (black) and SAT1 (blue) animals. Mean ± SEM. Two-way repeated measures ANOVA $F_{(1,19)} = 0.0056$, p = 0.94. (g) Spiking elicited by brief ChR2 activation (mean + SEM). ACC: 1.1 ± 0.5 (N = 5 mice, n = 13 cells); SAT1: 1.01 ± 0.04 (N = 3 mice, n = 8 cells). Two-tailed Mann-Whitney U test (U = 59.5, n = 13 and 8).

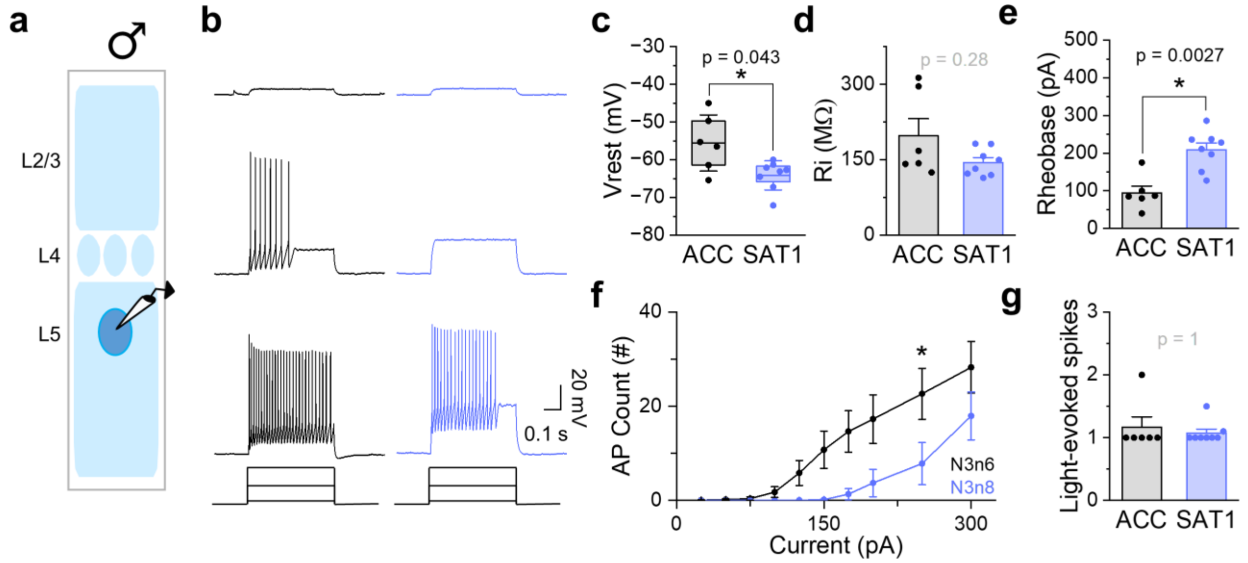


**Supplementary Fig. 6. Excitability of L5 PV neurons in males is reduced after training.**
(a) Schematic of the experiment setup. ChR2-expressing L5 PV neurons were targeted. (b) Representative firing of L5 fast-spiking PV neurons in response to 500 ms current injections (25, 150, 300 pA) after 1-2 days of acclimation (ACC: black) and 1 day of SAT (SAT1: blue). (c) Resting membrane potential (Vrest) comparison. ACC: -55.6 ± 7.5 mV (N = 3 mice, n = 6 cells); SAT1: -64.1 ± 3.9 mV (N = 3 mice, n = 8 cells). The box is the 25th and 75th quartiles, the whiskers are the SD, and the midline is the mean. Two-tailed Mann-Whitney U test (U = 40, n = 6 and 8). (d) Input resistance (Rin) comparison (mean + SEM). ACC: 197.6 ± 34.3 MΩ (N = 3 mice, n = 6 cells); SAT1: 144.7 ± 9.6 MΩ (N = 3 mice, n = 8 cells). Two-tailed Mann-Whitney U test (U = 33, n = 6 and 8). (e) Rheobase comparison (mean + SEM). ACC: 94.5 ± 18.1 pA (N = 3 mice, n = 6 cells); SAT1: 209.0 ± 18.0 pA (N = 3 mice, n = 8 cells). Two-tailed Mann-Whitney U test (U = 2, n = 6 and 8). (f) F-I curve of L5 PV neurons in ACC (black) and SAT1 (blue) animals. Mean ± SEM. Two-way repeated measures ANOVA $F_{(1,12)} = 6.7$, $p = 0.024$. *post hoc* Tukey's test. (g) Spiking elicited by brief ChR2 activation (mean + SEM). ACC: 1.2 ± 0.4 (N = 3 mice, n = 6 cells); SAT1: 1.1 ± 0.2 (N = 3 mice, n = 8 cells). Two-tailed Mann-Whitney U test (U = 23, n = 6 and 8). * $p < 0.05$.

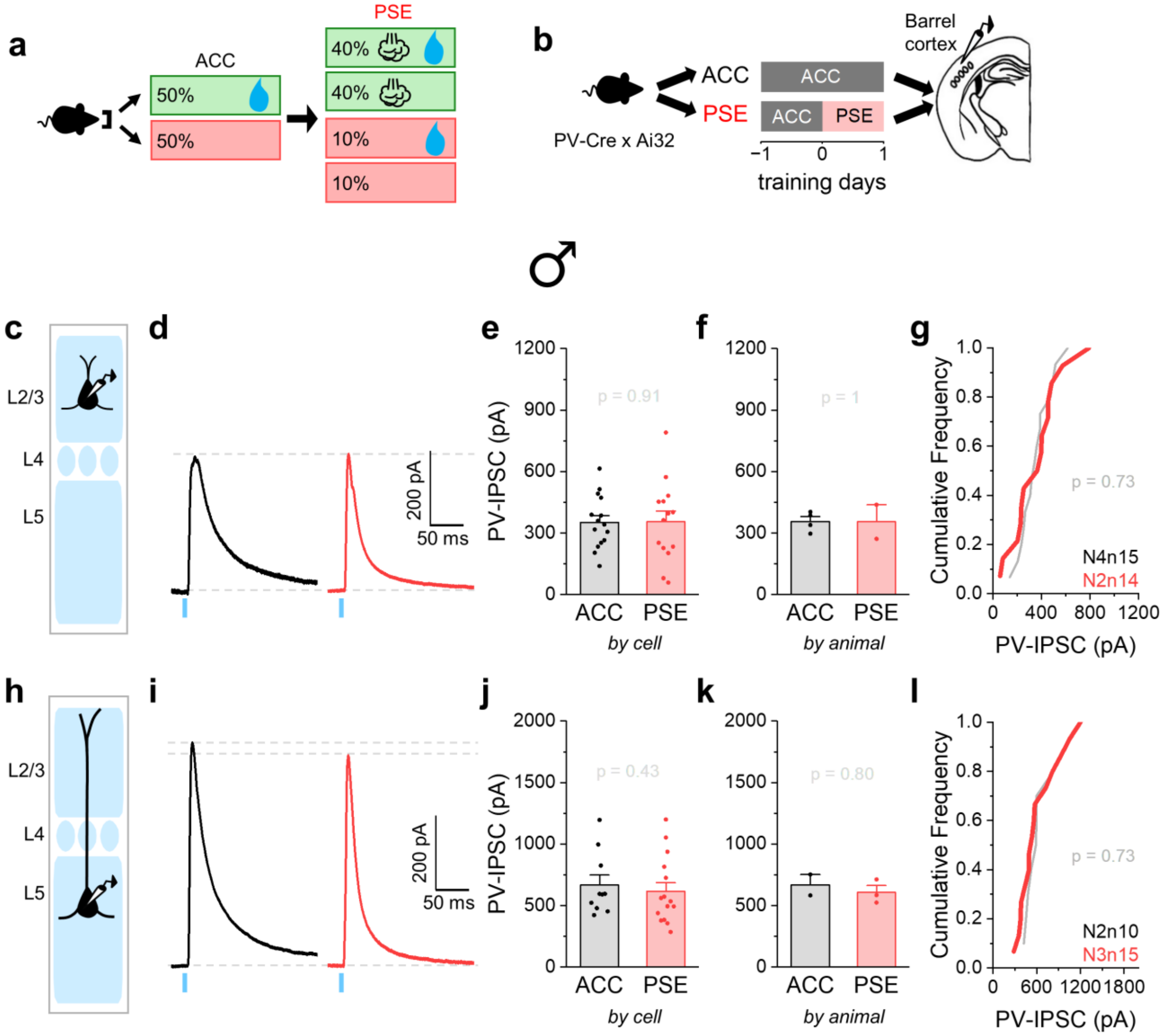


**Supplementary Fig. 7. Pseudotraining does not alter PV-mediated inhibition onto Pyr neurons in males.**

(a) Pseudotraining (PSE) paradigm. (b) Schematic of training paradigm and tissue preparation for PV-Cre x Ai32 mice. (c) Schematic of whole-cell recordings from L2/3 Pyr neurons. (d) Representative traces of ChR2-evoked PV-IPSCs (10 sweeps average) recorded from L2/3 Pyr neurons after 1-2 days of acclimation (ACC, black) and 1 day of PSE (red). (e) Peak PV-IPSC amplitudes in L2/3 Pyr neurons averaged across cells. ACC: 352.7 ± 33.4 pA (N = 4 mice, n = 15 cells); PSE: 355.1 ± 52.7 pA (N = 2 mice, n = 14 cells). Two-tailed Mann-Whitney U test (U = 108, n = 15 and 14). (f) Same as in (e), but averaged across animals. ACC: 356.6 ± 24.0 pA (N = 4 mice); PSE: 355.1 ± 83.3 pA (N = 2 mice). Two-tailed Mann-Whitney U test (U = 4, n = 4 and 2). (g) Cumulative distribution of PV-IPSC amplitudes in L2/3 Pyr neurons from ACC and PSE groups. Kolmogorov-Smirnov (K-S) test. (h-l) Same as in (c)-(g), but for L5 Pyr neurons. (j) ACC: 668.0 ± 81.1 pA (N = 2 mice, n = 10 cells); PSE: 615.7 ± 70.6 pA (N = 3 mice, n = 15 cells). Two-tailed Mann-Whitney U test (U = 90, n = 10 and 15). (k) ACC: 668.0 ± 85.1 pA (N = 2 mice); PSE: 607.6 ± 55.5 pA (N = 3 mice). Two-tailed Mann-Whitney U test (U = 4, n = 2 and 3). (l) K-S test. All bar graphs represent mean + SEM.

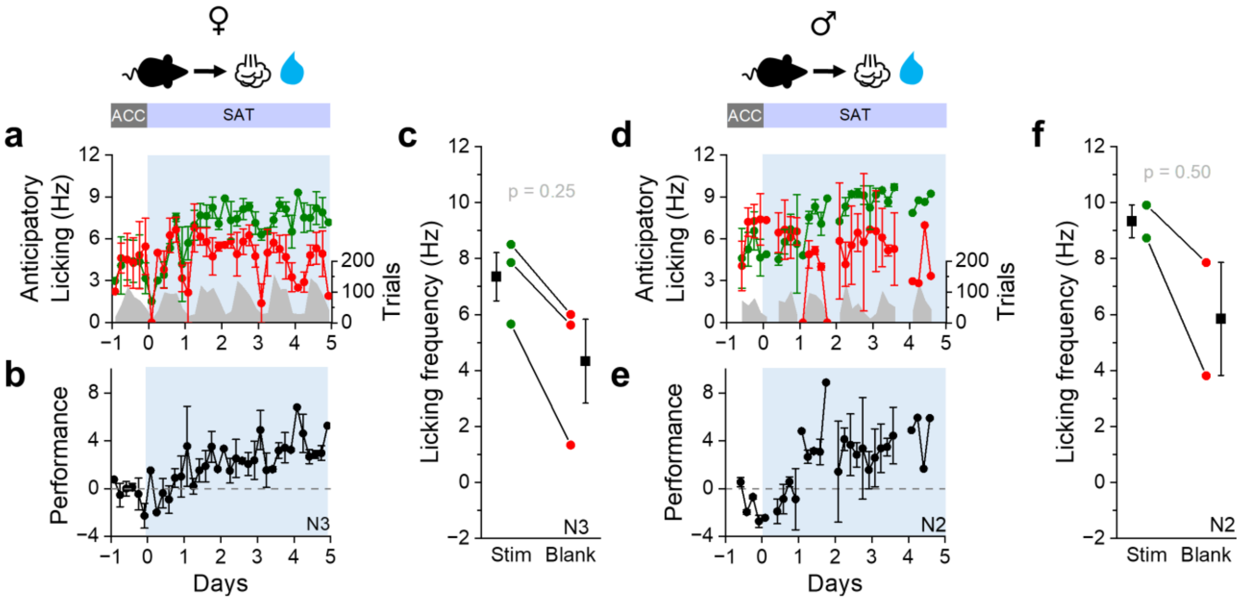


**Supplementary Fig. 8. Performance improves over time in both female and male mice during prolonged training.**

(a) Mean anticipatory licking frequency (Hz) for stimulus (green) and blank (red) trials of females following 5 days of SAT (SAT5). Blue shading indicates the training period. Gray shading indicates the distribution of average trial numbers across time bins. (b) Performance ($Lick_{stim}$ - $Lick_{blank}$; see Materials and Methods) was calculated for each 4-hr bin and averaged across animals. (c) Mean anticipatory licking frequency (Hz) during the last 20% of stimulus (green) and blank (red) trials for each female animal after SAT1. Black symbols represent group mean ± SEM. Stim: 7.3 ± 0.9 Hz; Blank: 4.3 ± 1.5 Hz (N = 3 mice). Paired-sample, Wilcoxon signed-rank test (Z = 1.34, n = 3). (d-f) Same as in (a)-(c), but for male animals. (f) Stim: 9.3 ± 0.6 Hz; Blank: 5.8 ± 2.0 Hz (N = 2 mice). Wilcoxon signed-rank test (*Z* = 0.89, n = 2).

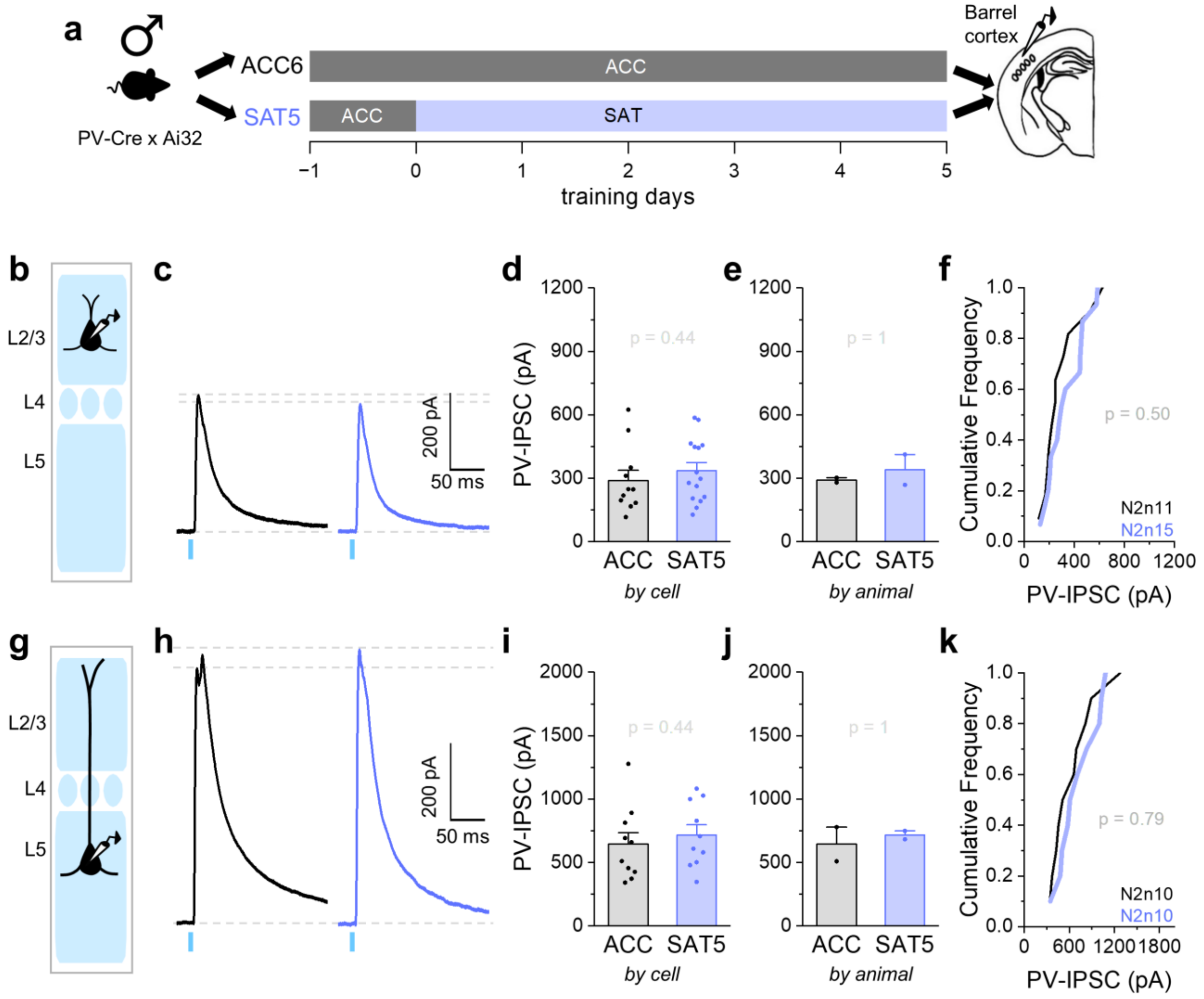


**Supplementary Fig. 9. Prolonged training does not alter PV-mediated inhibition onto Pyr neurons in males.**

(a) Schematic of training paradigm and tissue preparation for PV-Cre x Ai32 mice. (b) Schematic of whole-cell recordings from L2/3 Pyr neurons. (c) Representative traces of ChR2-evoked PV-IPSCs (10 sweeps average) recorded from L2/3 Pyr neurons after 6 days of acclimation (ACC, black) and 5 days of sensory association training (SAT5, blue). (d) Peak PV-IPSC amplitudes in L2/3 Pyr neurons averaged across cells. ACC: 290.2 ± 47.3 pA (N = 2 mice, n = 11 cells); SAT5: 355.9 ± 38.6 pA (N = 2 mice, n = 15 cells). Two-tailed Mann-Whitney U test (U = 67, n = 11 and 15). (e) Same as in (d), but averaged across animals. ACC: 291.2 ± 11.2 pA (N = 2 mice); SAT5: 340.7 ± 71.9 pA (N = 2 mice). Two-tailed Mann-Whitney U test (U = 2, n = 2 and 2). (f) Cumulative distribution of PV-IPSC amplitudes in L2/3 Pyr neurons from ACC and SAT5 groups. Kolmogorov-Smirnov (K-S) test. (g-k) Same as in (b)-(f), but for L5 Pyr neurons. (i) ACC: 644.4 ± 91.9 pA (N = 2 mice, n = 10 cells); SAT5: 716.4 ± 81.2 pA (N = 2 mice, n = 10 cells). Two-tailed Mann-Whitney U test (U = 39, n = 10 and 10). (j) ACC: 644.4 ± 134.6 pA (N = 2 mice); SAT5: 716.4 ± 32.8 pA (N = 2 mice). Two-tailed Mann-Whitney U test (U = 2, n = 2 and 2). (k) K-S test. All bar graphs represent mean + SEM.
